# Experimental evidence of male-male interaction in laboratory swarms of *Anopheles gambiae* mosquitoes

**DOI:** 10.1101/2025.11.06.686941

**Authors:** Gloria Iacomelli, Max Lombardi, Leonardo Parisi, Matteo Fiorini, Francesca Grucci, Alessio Lavorgna, Melania Ligato, Gaetano Zarcone, Matthew J. Peirce, Stefania Melillo, Roberta Spaccapelo

## Abstract

Mosquitoes mostly mate in the context of swarms: to facilitate encounters with females, males form disordered aggregations over a visual marker, which serves as a positional reference. Due to the scarcity of high-resolution data, it remains unclear whether swarms are the result of individuals interacting independently with the marker, or whether they are the expression of a genuine collective behavior produced by spontaneous interaction between individuals. Here, with a unique dataset comprising three–dimensional trajectories of 30 laboratory swarms of different size (ranging from 80 to 400 mosquitoes), we investigate swarming behavior of *Anopheles gambiae* mosquitoes. With a statistical physics approach, we provide experimental evidence of an effective male-male interaction. We find that individual speed fluctuations are strongly correlated in space, meaning that mosquitoes in close proximity tend to display similar deviations from the group average, effectively flying at a similar speed. We prove that this correlation is not compatible with a random arrangement of individuals, indicating that an interaction process is at play.

## 1 Introduction

Collective behavior is a widespread phenomenon, observed across many biological systems, from cells in a colony to vertebrates in a herd. It emerges when a large number of individuals self-organize and, through mutual interaction, exchange information that can be transferred to the entire group. In its most fascinating manifestations, interaction leads to global order. This is the case, among others, with the synchronized flashing of fireflies [1, 2], or the coordinated motion of birds and fish, [3–6].

While there is general consensus [7, 8] that order arises from the interaction between individuals, the debate on the nature of the behavior in disordered systems is still active, with insect swarms serving as primary examples [9–11]. Swarms are indeed the paradigm of disordered systems: hundreds of insects - exclusively males – congregate about a common marker [12], to facilitate encounters with females, who will join the group only briefly to mate [13]. As a whole, the swarm maintains a stable position over the marker, which is used as a positional reference [14]. But, at an individual level, insects in the swarm follow apparently erratic flight patterns.

The lack of global order, combined with the presence of a marker, supports two competing interpretations. In one, the visual attraction towards the marker dominates - if not completely determines - the insects’ dynamics [9, 15]. In the other, the marker plays a more marginal role, and the interaction between individuals gives rise to collective behavior [11].

In swarms of non-biting midges (family Chironomidae), the emergence of an authentic collective behavior has been proved through field experimental data [11, 16, 17], and confirmed by theory and simulations [18]. In mosquitoes, instead, there are still no conclusive studies. Research is mainly focused on the characterization of individual dynamics and overall swarm structure [19–21], as well as on the mechanisms of courtship and mating [22–24], with only a very few studies specifically addressing male-male interaction, [25–27]. Three dimensional field data on small swarms (less than 100 individuals) of *Anopheles gambiae* and *Anopheles coluzzii* [25, 26] suggest the presence of a short-range interaction, which leads to alignment in the direction of flight. A different form of interaction, mediated by the mosquito auditory system, has been proposed in *Aedes aegypti* [27], where the emergence of acoustic order is observed in short arrays (up to seven individuals) of tethered mosquitoes.

Here, with a unique dataset of 30 swarms, ranging from 80 to 400 individuals, we investigate lab–based swarming behavior in a lab-adapted strain (G3) of *Anopheles gambiae* mosquitoes. Using a statistical physics approach, we provide experimental evidence of a male-male interaction that, unlike previous studies [11, 25, 26], does not act on mosquitoes’ direction of flight, but rather produces a long-range correlation of their speed.

## 2 Results

### 2.1 Experiments and data

We perform laboratory experiments on mosquitoes, *Anopheles gambiae* (strain G3), swarming in a large cage (a room 5.*0m* × 3.5*m* × 2.6*m*), where we artificially reproduce visual stimuli needed for mosquitoes to swarm, see Methods and [20].

Data are collected with a 3*D* camera system: three synchronized high-speed cameras (IDT-M5) recording the same swarm from different positions, so that in post-processing, with the computer vision software GReTA [28], we are able to reconstruct individual trajectories of each individual in the swarm.

Data presented in the main section of this work comprise 30 swarms, ranging from 80 to 400 individuals. A distinct dataset (41 smaller swarms ranging from 10 to 80 individuals) is included in SI, to highlight the relevance of swarm size for behavioral analysis. Data belong to two different experimental campaigns: the first conducted in 2020/2021 with cameras shooting at 170fps and acquisitions lasting ∼ 15*s*, the second conducted in 2024 at 80fps with acquisitions lasting ∼ 30*s*.

### 2.2 Localization within the swarm

The fundamental assumption underpinning most advanced models for swarming dynamics [29], and more generally of statistical physics, is that individuals within a group are indistinguishable. At a given instant in time they may be located in different positions and display different kinematic attributes, but these differences are solely due to statistical fluctuations, and essentially vanish when observing the system over time. In contrast, we find that, despite being part of a swarm, mosquitoes maintain their individuality and exhibit specific characteristics that are not compatible with purely statistical fluctuations.

The analysis of individual trajectories shows that each mosquito performs consecutive loops, [20], around a specific position, slightly different from that of the other members of the group. In performing these loops, it explores a small portion of the volume occupied by the entire swarm, see Fig.1. Similarly, we find that each mosquito flies at its own characteristic speed, and it samples only a narrow range of the speeds observed at a group level, see Fig.2.

**Fig. 1.**
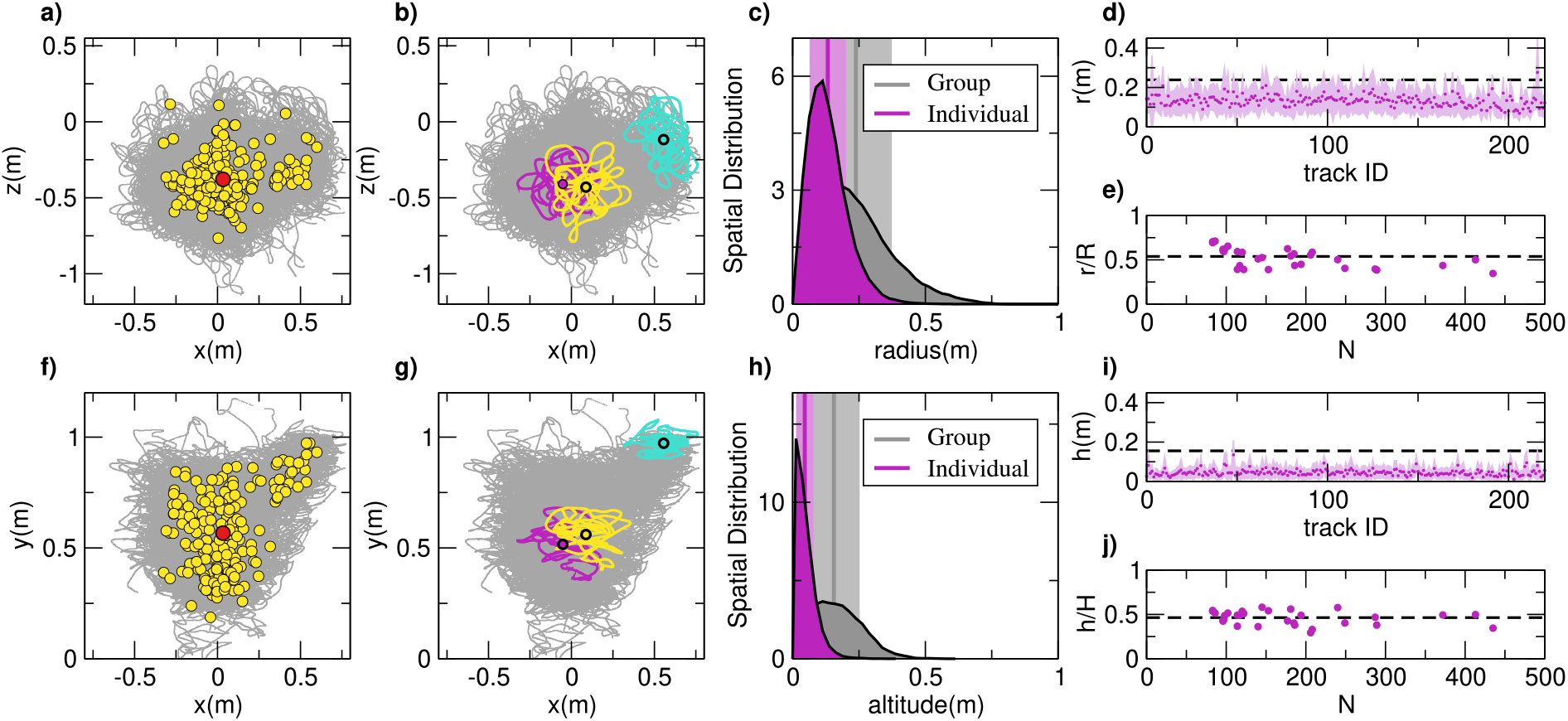
Spatial localization. Swarm 20200928_*ACQ*3. Number of mosquitoes ∼ 200, duration 2700 frames at 170fps, corresponding to 15.88s. **a**. Top view (*xz*-plane). In dark gray the 3*D* reconstruction of the swarm. The red circle represents the swarm centroid. The yellow circles indicate individual centroids, one for each mosquito in the swarm. **b**. Top view, *xz*– plane. In dark gray the 3*D* reconstruction of the swarm. In turquoise, yellow and purple three different trajectories highlight the spatial individual localization. Mosquitoes do not explore the entire volume occupied by the swarm, but only a small portion of it. The three black circles represent the individual centroids of the three trajectories. **c**. On the horizontal plane, the group spatial distribution (dark gray) is much broader than the individual spatial distribution (purple). Group radius is equal to *R* = 0.24 ± 0.14m, the individual radius *r* = 0.13 ± 0.07m. The vertical lines and vertical shaded areas represent the mean and standard deviation, gray for the group and purple for individuals. On the right of the two distributions, a schematic description of the two variables *r* (indicated by white within purple circle) and *R* (indicated by black within gray circle), where the large dark gray circle represents the swarm and the three small circles the volumes occupied by the three trajectories highlighted in yellow, purple and turquoise in panel b. **d**. On the horizontal plane, individual radii show low variability within the trajectories of the swarm, and are consistently smaller than the group radius (the black dashed line.) **e**. On the horizontal plane, the ratio between individual and group radii, *r/R*, is approximately equal to 0.5 (the black dashed line) in all the 30 swarms of the dataset, regardless of the number of mosquitoes in the swarm, *N*. **f**. Side view (*xy*-plane). In dark gray the 3*D* reconstruction of the swarm. The red circle represents the swarm centroid. The yellow circles indicate individual centroids, one for each mosquito in the swarm. **g**. Side view. In dark gray the 3*D* reconstruction of the swarm. In turquoise, yellow and purple three different trajectories, the same ones shown in panel b, highlight the spatial individual localization. **h**. Along the vertical axis, the group spatial distribution (dark gray) is much broader than the individual spatial distribution (purple). Group altitude is equal to *H* = 0.16 ± 0.10m, the individual altitude *h* = 0.05 ± 0.03m. The vertical lines and vertical areas represent the mean and standard deviation, gray for the group and purple for individuals. On the right of the two distributions, a schematic description of the two variables *h* (indicated by white within purple circle) and *H* (indicated by black within gray circle), where the large dark gray rectangle represents the swarm and the three small rectangles the volumes occupied by the three trajectories highlighted in yellow, purple and turquoise in panel g. **i**. Along the vertical axis, individual altitudes show low variability within the trajectories of the swarm, and they are consistently smaller than the group altitude (the black dashed line.) **j**. Along the vertical axis, the ratio between individual and group altitudes, *h/H*, is approximately equal to 0.5 (the black dashed line) in the 30 swarms of the dataset, regardless of the number of mosquitoes in the swarm, *N*.

**Fig. 2.**
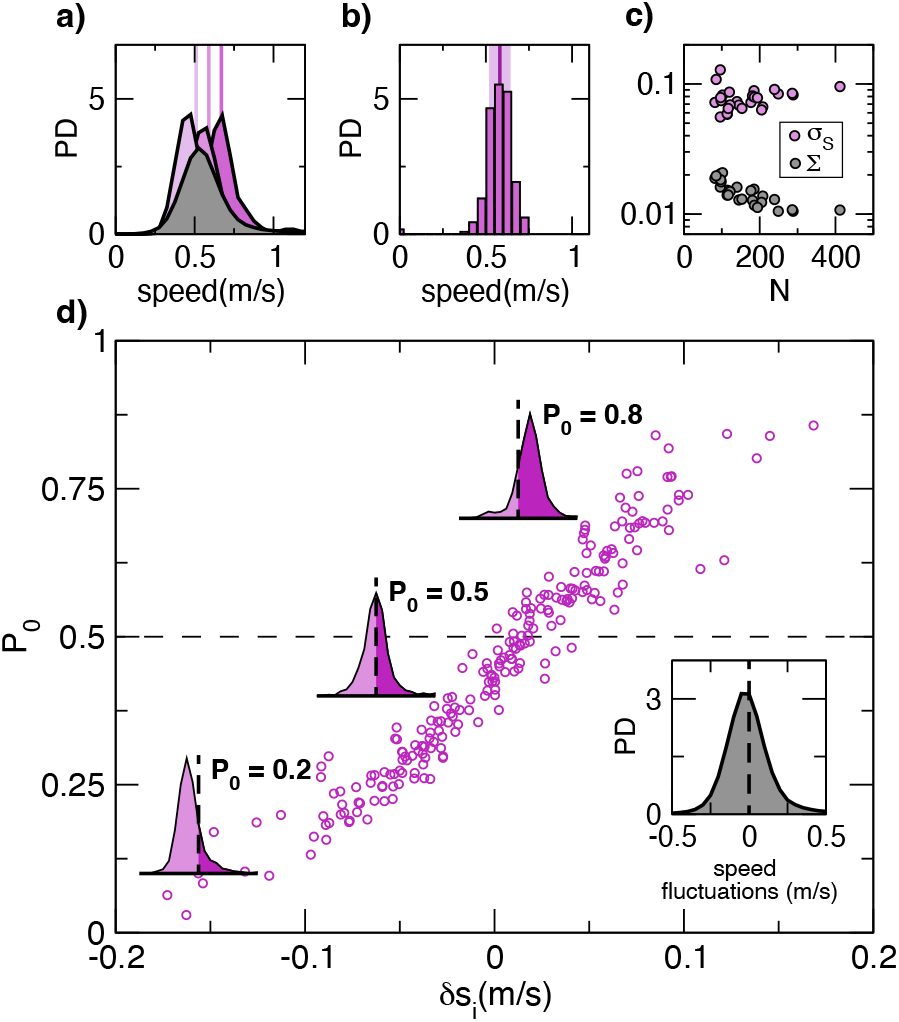
Speed localization. a. In dark gray a typical group probability distribution. In purple, individual probability distributions of three different mosquitoes, whose mean speeds are highlighted with the purple vertical lines. **b**. The probability distribution of the individual mean speeds for the 200 mosquitoes belonging to the swarm 20200928 *ACQ*3. Mean speeds are in the range between 0.37m/s and 0.75m/s. This distribution displays large fluctuations, with standard deviation equal to 0.06m/s, highlighted with the light purple stripe. **c**. The fluctuations of individual mean speeds measured from the data (*σ*_*S*_, in purple) and estimated from the group probability distribution (Σ, in gray) for all the swarms of our dataset, as a function of the size of the system. Measured standard deviations are consistently greater, up to ten times larger, than their expected values in the absence of localization. **d**. For the same swarm shown in panels a and b, the probability, *P*_0_, that individual speed fluctuations are positive as a function of *δs*_*i*_. *P*_0_ spans the entire range from 0.1 to 0.9, confirming speed localization in mosquitoes. In the absence of localization, *P*_0_ is equal to 0.5, the dashed horizontal line in the plot. As expected, *P*_0_ increases with increasing *δs*_*i*_. In the inset: the group probability distribution of the fluctuations.

This leads to both spatial and speed *localization*.

#### Spatial localization

We give a quantitative description of the spatial localization, by estimating and comparing the extent of the swarm as a whole and that of individual trajectories.

To this aim, we compute the swarm centroid (the red circle in Fig.1a,f), averaging over the positions of all mosquitoes at all instants of time. We then measure the distance of each mosquito from the swarm centroid, and we define the *group* spatial distribution as the probability distribution of these distances, see Methods. This spatial distribution describes how frequently an individual is found at a given distance from the swarm center. Its width provides a rough estimate of the spread of the swarm, while its mean is a measure of the swarm size.

In a similar way, we compute the centroid of each trajectory (the yellow circles in Fig.1a,f - one for each mosquito - and in the black circles in Fig.1b,g). We then measure the distance of each mosquito from its *own* centroid, and we define the *individual* spatial distribution as the probability distribution of these distances, aggregating all mosquitoes in the swarm, see Methods. The width of the individual spatial distribution gives a rough estimate of the typical spread of single trajectories, while its mean is a measure of the size of the region explored by each individual.

Mosquitoes move predominantly on the horizontal plane (*xz*-plane), with limited changes in altitude (*y*-axis), [20]. Because of this anisotropy, we measure group and individual spatial extents on the horizontal plane (radius) separately from the elongation along the vertical axis (altitude).

On the horizontal plane, the group spatial distribution is significantly broader than the individual one, see Fig.1c. For the swarm shown in Fig.1, the group radius is *R* = 0.24 ± 0.13m, while the individual one is *r* = 0.13 ± 0.07m, with very low variability between mosquitoes, see Fig.1d. This results in a ratio *r/R* ∼ 0.5. Similarly, along the vertical axis, the swarm is much more elongated than individual trajectories. The typical vertical extent for the swarm in Fig.1h is equal to *H* = 0.16 ± 0.10m, while the individual elongation is *h* = 0.05 ± 0.03m, again with low variability between different mosquitoes and with a ratio *h/H* ∼ 0.5, Fig.1i.

By performing this same analysis over all the swarms of our dataset, we find that *r/R* and *h/H* are consistently close to 0.5, see Fig.1e,j. This confirms that mosquitoes do not explore the entire volume occupied by the swarm, but instead remain localized in a small region, approximately one eighth of the total volume.

#### Speed localization

Through the analysis of individual speed, we find a phenomenon that, by analogy to what we have already observed for mosquito positions, we refer to as *speed localization*: individual mosquitoes do not explore the full range of speed covered by the entire swarm, but only a portion of it.

A first hint of speed localization emerges from the comparison between individual and group probability distributions, see Fig.2a, where we show in dark gray the group distribution and, in purple, individual distributions for three different mosquitoes. In the absence of localization, we would expect the group and individuals to share the same statistical properties. All individuals would display a similar speed over time, so that their probability distributions would be identical and coincide with that of the group. Under this assumption, variations in individual mean speeds whose distribution is presented in Fig.2b, would reflect only statistical fluctuations and could be estimated by the standard error of the group distribution, Σ, see Methods. Having access to full trajectories, we are also able to measure these variations directly from the data, in the form of the standard deviation of individual mean speeds, *σ*_*S*_, see Methods. We find *σ*_*S*_ in the interval between 0.06m/s and 0.13m/s, with Σ in the interval between 0.01m/s and 0.02m/s, see Fig.2c. The comparison of these two quantities indicate that in our swarms the variation of the mean speed across individuals cannot be explained by statistical fluctuations alone.

The effect of speed localization becomes clearer when shifting our focus from absolute speeds to deviations from the group average. In particular, the speed fluctuation of mosquito *i* at time *t* is defined as:

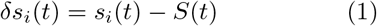

where *S*(*t*) represents the group average speed at time *t, s*_*i*_(*t*) the speed of mosquito *i* at time *t*, and *δs*_*i*_(*t*) its fluctuation. This transformation moves us from speed — always a positive quantity — to its fluctuation, which can be positive or negative. If a mosquito is flying faster than the group average, the fluctuation is positive; if slower, it is negative.

At a group level, the probability distribution of the fluctuations is centered around 0m/s, and it is nearly symmetric, see the inset of Fig.2d: At any one time, there is approximately a 50% chance of finding a positive fluctuation (a mosquito flying faster than the group average), and a 50% chance of a negative fluctuation (a mosquito flying slower than the group). But when we examine speed fluctuations at an individual level, this symmetry breaks down. Individual distributions are not centered around zero, but they are unbalanced toward positive or negative fluctuations.

This individual asymmetry is the hallmark of speed localization. We quantify the asymmetry by computing, for each mosquito, the probability, *P*_0_, that its speed fluctuations are positive. We essentially evaluate the proportion of time (relative to each trajectory’s duration) that each mosquito spends flying faster than the group average. This corresponds to the area, highlighted in dark purple in the histograms in Fig.2d, under the distribution to the right of the *y*-axis, see Methods. In the absence of localization, *P*_0_ should be approximately 0.5. We find instead that *P*_0_ spans the full range from 0.1 - mosquitoes that fly slower than the group average for most of the time - to 0.9 - mosquitoes that fly faster than the group average for most of the time. This wide variation confirms the presence of speed localization in our swarms.

### 2.3 Experimental evidence of male-male interaction

In biological systems, interaction is typically driven by sensory processes, involving vision, hearing, and other senses [30–33]. Individuals perceive the other members of the group, and adjust their own behavior, synchronizing their state. The physical limit of sensory perception defines the *interaction range*, which may be quite short (in mosquitoes hearing range is of the order of 10-20cm [34, 35]). Despite the limited interaction range, information can travel much greater distances, thanks to a sort of *word of mouth* mechanism, see Fig.3a: *A* interacts with *B*, and *B* also interacts with *C*, who lies outside *A*’s interaction range. By interacting, *A* and *B* adjust their behavior, as do *B* and *C*. As a result, the states of the three individuals are synchronized. Thanks to the indirect exchange of information between *A* and *C*, mediated by *B*, the system exhibits correlation over a distance (that between *A* and *C*) that exceeds the interaction range. In physics, this indirect synchronization, produced by a chain of interactions, is referred to as correlation.

**Fig. 3.**
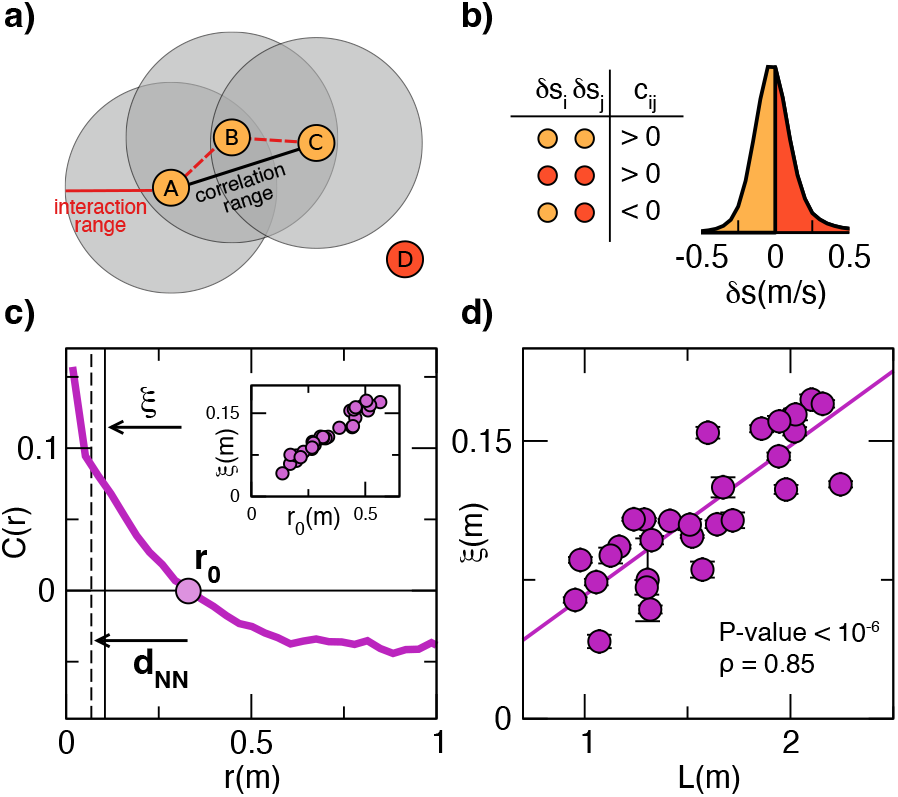
Speed correlation. a. Interaction vs correlation. Interaction and correlation are both related to a transfer of information. Despite the short range of interaction, information can travel greater distances: *A* interacts with *B. B* interacts also with *C*, which exceeds A’s interaction range. Due to the interaction, *A* and *B* synchronize their speed, so do *B* and *C*. As a result, the three particles are synchronized, hence correlated. The correlation range of the system is defined by the distance between *A* and *C*, that is larger than the interaction range. *D* exceeds the interaction ranges of the other particles. **b**. The group distribution of the speed fluctuations, with light and dark orange differentiating between negative and positive fluctuations. Two particles having same sign of fluctuations produce a positive correlation coefficient *c*_*ij*_. **c**. The correlation function has a non–trivial trend. It is positive for short distances. It decreases at increasing distances, and crosses the *x*–axis at *r*_0_, which defines the correlation range. *r*_0_ is much larger than the nearest neighbor distance, *d*_*NN*_. The correlation length, *ξ* is computed following [4]. Inset: correlation length, *ξ*, scales linearly with the correlation range (*ρ* = 0.99, p-value < 10^−6^). **d**. Across all swarms of the dataset, the correlation length, *ξ*, scales linearly with the size of the system, *L* (*ρ* = 0.85, p-value < 10^−6^). In the plot, bars are standard deviations. This linear trend represents the finite size expression of the scale–free property of the correlation function: there is no typical correlation length for our swarms; but the only relevant scale for the system is its size.

Interaction and correlation are distinct, yet closely related, concepts. They both refer to an exchange of information that is direct in the case of interaction, but indirect in the case of correlation. Measuring how mosquitoes react to each other is relatively simple with tethered mosquitoes [27, 36]. However, this kind of measurement is much more complicated in free–flying swarms. In these specific situations, where a direct measure of interaction is not feasible, correlation functions provide a powerful tool to reveal the presence of interaction within the system.

#### Spatial correlation functions of speed fluctuations

We measure whether and to what extent (in space), mosquitoes tend to synchronize deviations in their speed from the group average, by computing the spatial correlation function of the speed, in its connected declination, [11, 37]. It is worth noting that, unlike previous studies on insect swarms [11, 16], we investigate the correlation function of the speed, which does not refer to the direction of flight, as the velocity does. As shown in SI, our swarms display no significant correlation of velocity.

We quantify the synchronization between two mosquitoes, by computing their correlation coefficient:

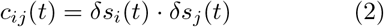

where *δs*_*i*_(*t*), *δs*_*j*_(*t*), defined as in eq.(1), denote the speed fluctuations of individuals *i* and *j* at time *t*. This product is positive if fluctuations are both positive or both negative (both mosquitoes flying faster or both fly slower than the group average), see Fig.3b. Conversely, it is negative if one is positive and the other is negative (one mosquito flying faster than the group average, the other flying slower). Therefore, a positive *c*_*ij*_ indicates a tendency towards synchronization.

We can now group pairs of mosquitoes based on their mutual distance, and compute the average of the correlation coefficient over each bin. With this procedure we associate a correlation value with each distance, thereby defining the spatial correlation function, *C*(*r*) (see Methods for a formal definition). The *C*(*r*) essentially defines what is the typical correlation coefficient of two mosquitoes at a mutual distance *r*. Across all swarms of our dataset, it exhibits a characteristic trend: it is positive for small values of *r*, meaning that mosquitoes in close proximity display similar speed fluctuations. It then decays with increasing distance, crossing the *x*–axis at a distance *r*_0_, see Fig.3c, which defines the correlation range and gives a rough estimate of the size of the correlated domains.

We also give an estimate of the size of the correlated domains as the standard correlation length, *ξ*, that we compute using the integral form in [4]. This definition of *ξ* is general and robust, since it takes into account the entire shape of the correlation function up to *r*_0_. In our case, across all swarms, *ξ* grows linearly with *r*_0_, see inset in Fig.3c, varies in the range between 0.04m and 0.16m (see Table S1) and it scales linearly (Pearson coefficient *ρ* = 0.85, *p* − *Value* < 10^−6^) with the size of the swarm, *L*, see Fig.3d. This linear scaling is the finite-size expression of what is known in physics as *scale-free* correlation, a property also observed in other biological systems [4, 11]. It implies that there is no characteristic distance beyond which information transfer becomes negligible. Rather, the only relevant scale is the system size itself — meaning that information can propagate across the entire system, regardless of its size.

#### Evidence of interaction

In the absence of spatial and speed localization, non–trivial and scale– free spatial correlation would suggest that an *effective* interaction, acting on the speed domain, is at work in swarms of mosquitoes. But this correlation inherently involves both space and speed. In a system such as ours, which exhibits localization, it is then natural to wonder whether correlation is the genuine expression of interaction, or whether it is merely an epiphenomenon of the system properties.

To discriminate between these two scenarios, we first investigate the spatial dependence of the speed within our swarms. We then test our data against null cases, consisting of systems of non-interacting particles that do exhibit spatial and speed localization. In doing this, we prove that, while such localization contributes to the correlation, it cannot account for its full magnitude. Instead, correlation is primarily driven by the instantaneous spatial arrangement of the mosquitoes and by the synchronization of their speed fluctuations.

#### Lack of speed spatial structure

The correlation we find in our swarms implies that mosquitoes in close proximity exhibit similar speeds. Without invoking a speed–based interaction, we might explain correlation with the more parsimonious assumption that the speed of each individual depends on its spatial position. Under this mechanism, mosquitoes occupying the same region of the swarm would be subject to similar local conditions and, for this reason, display similar speeds. Correlation could thus reflect the spatial context, rather than a direct synchronization between individuals. To test this hypothesis, we evaluate the dependence of individual speed fluctuations on their distance from the swarm center (see SI), as suggested by the model in [38]. We find no significant trend, with Pearson’s coefficient in the range between − 0.04 and 0.06; the same holds with respect to the three spatial coordinates *x, y*, and *z*.

The position dependence of the speed does not necessarily manifest as a measurable trend. We therefore extend our analysis to investigate whether our swarms can be partitioned into speed-based regions, namely clusters of mosquitoes flying at similar speeds, distinct from those in nearby spatial regions. To this end, we apply a connected component clustering technique. Two mosquitoes are linked if the difference between their speed fluctuations is below a specific threshold (see Methods and Fig.4a). Clusters are then defined as groups of mosquitoes that are directly or indirectly connected through these links, meaning that any two individuals in the same cluster can be reached through a chain of such connections. Because of this chain mechanism, pairs of mosquitoes within the same cluster may exhibit speed fluctuations larger than the threshold, provided they are connected through an intermediate sequence of links. In contrast, mosquitoes within distinct clusters are separated by a gap in the speed fluctuations that cannot be bridged by any intermediate individual, see Fig.4e.

**Fig. 4.**
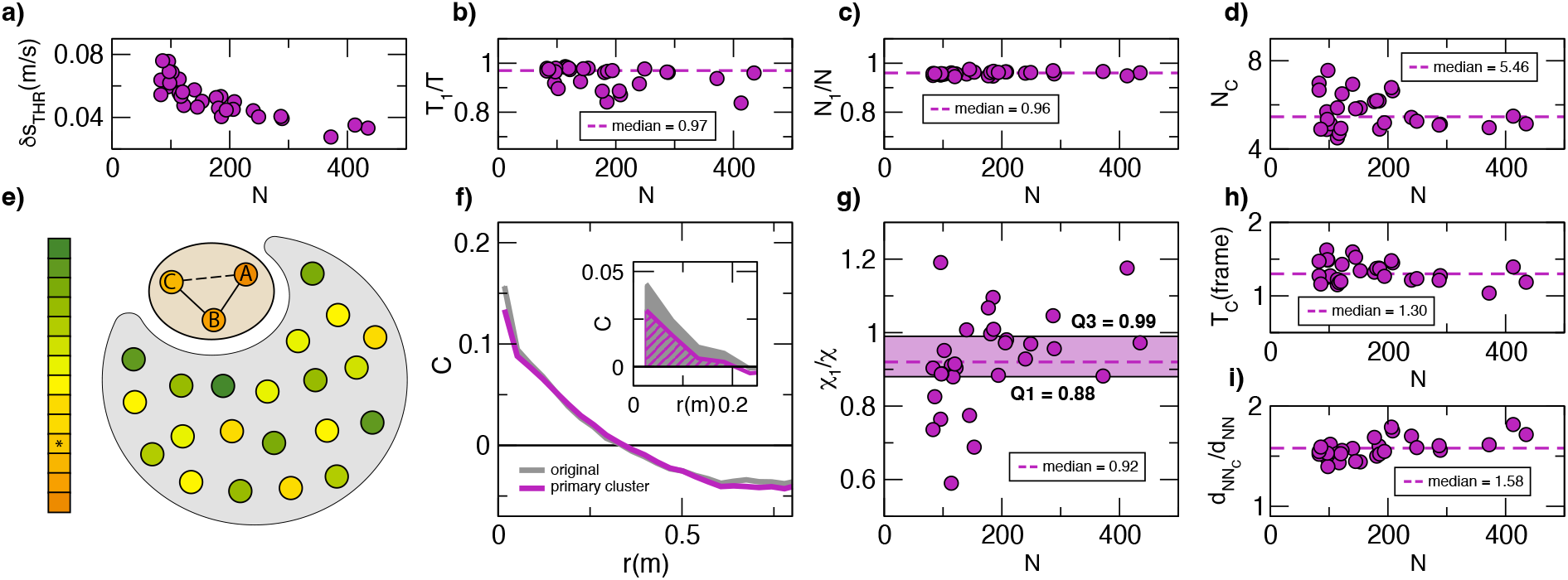
Lack of speed spatial structure. a. The threshold used in the speed clustering, *δs*_*T HR*_, as a function of the number of mosquitoes in the swarm, *N*. *δs*_*T HR*_ is defined through the elbow procedure, see Methods. It decreases with *N*, being in the range between 0.004m/s and 0.008m/s; **b**. The ratio between the number of frames with a single cluster, *T*_1_, and the acquisition duration, *T*, as a function of the number of mosquitoes, *N*, for all our swarms. Median value is equal to 0.97, with *T*_1_*/T* > 0.8 in all our swarms; **c**. The primary cluster is about the same size as the swarm. The ratio between the number of mosquitoes in the primary cluster, *N*_1_, and *N* is always larger than 0.9, with typical value equal to 0.96; **d**. The mean number of mosquitoes in the secondary clusters, *N*_*C*_, as a function of the number of mosquitoes in the swarm, *N*. Secondary clusters are consistently small, with a median size of 5.46 and always fewer than 8 individuals; **e**. A schematic illustration of how mosquitoes are clustered based on their speed. Dots represent individual mosquitoes, while their color encodes their speed (slow mosquitoes are shown in orange, while in green the fast ones). Mosquitoes with speed fluctuation differences below a given threshold are connected. In the example, each mosquito can only be linked to others whose colors fall within one color interval of the palette. Mosquito A is directly connected to mosquito B (solid black line), and B is in turn connected to mosquito C. As a result, A and C are indirectly connected (dashed black line) through B. The three mosquitoes A, B, and C therefore belong to the same cluster, highlighted by the light orange background. All other mosquitoes, colored from yellow to green, form a separate cluster, highlighted in gray. Because of the chain of links, mosquitoes in the same clusters may exhibit speed fluctuations whose difference is above the threshold, as for instance yellow and green dots in the gray cluster. The two clusters, the primary (highlighted in gray) and the secondary (light orange), are separated and cannot be connected because of a missing color, the one highlighted with a star in the palette, producing a gap in the speed intervals; **f**. The comparison between the swarm correlation (dark gray) and the correlation of the primary cluster (purple), for the swarm in acquisition 20200928 *ACQ*3, shows that the primary cluster drives the correlation of the group. **Inset:** comparison for the swarm with the smallest value of *χ*_1_*/χ* (see panel f). The shaded dark gray and hatched purple areas under the curves represent the susceptibilities *χ* (swarm) and *χ*_1_ (primary cluster), respectively. **g**. The ratio *χ*_1_*/χ* as a function of *N*. *χ*_1_*/χ* is always larger than 0.6, having median equal to 0.92 (first quartile 0.88, third quartile 0.99), indicating that the primary cluster largely drives the correlation function. For some of our swarms, *χ*_1_ exceeds *χ* (*χ*_1_*/χ* > 1), suggesting that in those cases secondary clusters tend to reduce the overall correlation. **h**. The temporal persistence of secondary clusters as a function of *N*. Persistence is defined, for each swarm, as the time-average number of consecutive frames in which we find more than one cluster. The median value is 1.3, indicating that clusters form and disrupt very quickly; **i**. The mean over time of the ratio between the distance to the nearest neighbor within the secondary clusters and within the whole swarm, as a function of *N*. This ratio is always greater than 1 (median 1.56), indicating that secondary clusters are less dense than the swarm;

For most of the time, our swarms consist of a single cluster, as large as the entire swarm, see Fig.4b-c. In the remaining frames, we identify a primary cluster along with a few secondary clusters that are significantly smaller, present a lower spatial density than that of the primary one, and exhibit a short temporal persistence (Fig.4d-h-i). These characteristics indicate that correlation is not driven by a specific spatial structure of the speed, which is indeed lacking in our swarms. As a further confirmation, we compute the correlation function restricted to the primary cluster (Fig.4f), and its susceptibility, *χ*_*P*_, as the integral of the correlation up to *r*_0_, see Methods. Similarly, we define *χ* as the susceptibility of the entire swarm. The ratio *χ*_*P*_ */χ*, with median value equal to 0.92 and consistently larger than 0.6 as shown in Fig.4g, confirms that the primary cluster provides the dominant contribution to the correlation. Overall, despite the existence of these secondary clusters, mosquitoes in our swarms do not show a spatial segregation based on speed; instead their speed fluctuations are well mixed in space, while remaining long range correlated.

#### Relevance of spatial arrangement

With the aim of evaluating the relevance of the specific spatial arrangement of mosquitoes in the swarm, we generate artificial swarms directly from the experimental data. Specifically, we disrupt potential interaction links through a shuffling procedure applied to the mosquitoes’ trajectory centroids, see Fig.5a for an example and Methods for formal details. With this procedure, artificial swarms present the same localization parameters, and the same speed probability distributions as real swarms, from which they differ only in the mosquitoes’ positional arrangement.

**Fig. 5.**
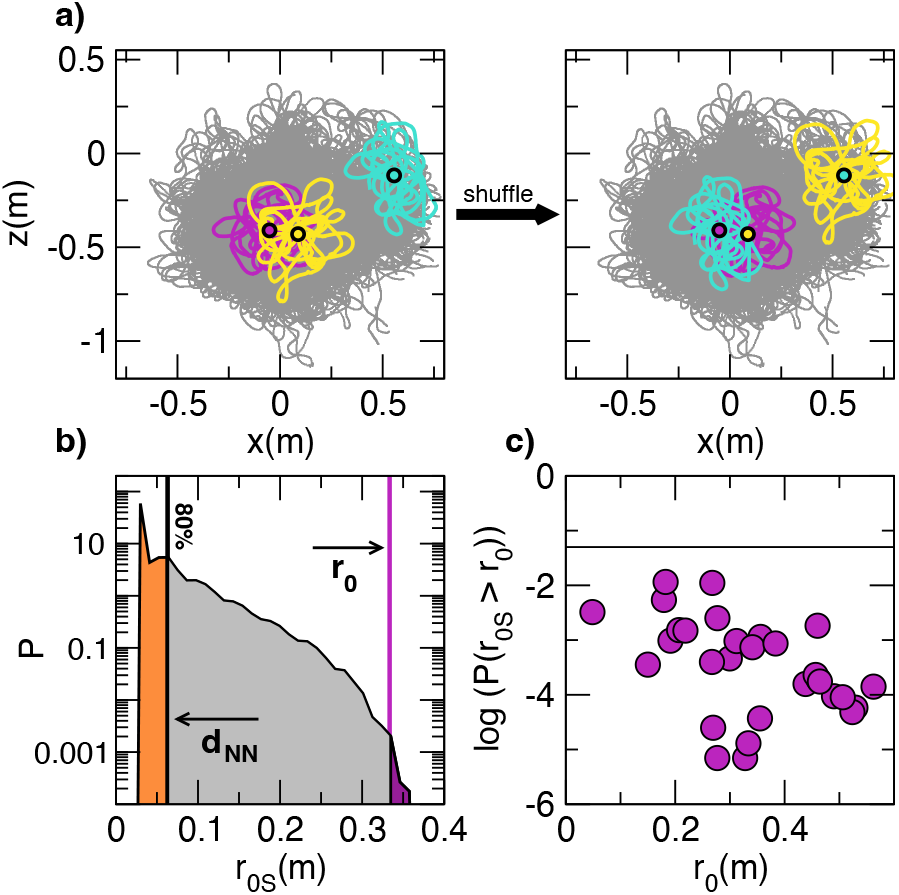
Relevance of spatial arrangement. a. We generate artificial swarms of non–interacting particles, that present the same localization properties and speed distributions of real swarms. We shift mosquito positions following a shuffling procedure on mosquito centroids. In the example, the centroid of the turquoise trajectory is assigned to the yellow mosquito, which is then shifted from the center of the swarm to the upper right corner; the centroids of the yellow and purple trajectories are assigned to the purple and turquoise mosquitoes respectively. The yellow and purple mosquitoes, originally close and potentially interacting, are located in two distant regions of the shuffled swarm, and thus no longer interact. **b**. The probability distribution of the correlation range for the 10^6^ correlation functions of the shuffles swarms. In 80% of the cases, the system presents a negligible correlation, with a correlation range smaller than the nearest neighbor distance, *d*_*NN*_. The area (highlighted in purple) below the distribution, on the right of the correlation range of the original real swarm, represents the probability of finding in the artificial swarm a correlation range larger than that of the original swarm. **c**. The logarithm of the probability *P* (*r*_0*S*_ > *r*_0_) for all the swarms of the dataset. Probability values are between 7 · 10^−6^ and 1 · 10^−2^, always smaller than the p–value threshold for significance of 0.05 (highlighted with the horizontal line in the plot). This suggests that the correlation range observed in real swarms is not compatible with a random arrangement of mosquitoes, providing evidence that the observed correlations are not merely a consequence of localization, and thus strongly indicate male–male interaction in our swarms.

For each real swarm, we generate 10^6^ shuffled instances, and we compute their correlation ranges, *r*_0*S*_. The probability distribution of *r*_0*S*_, in Fig.5b, shows that in most of the cases *r*_0*S*_ is smaller than the distance to the nearest neighbor, *d*_*NN*_, indicating that the correlation is negligible. From this same distribution we measure the probability *P*(*r*_0*S*_ ≥ *r*_0_) that the correlation range in shuffled swarms is equal or larger than that of real ones, the purple area in Fig.5b. In doing so, we essentially compute the p–value for the correlation range. Across our entire dataset, we find p-values between 7 ·10^−6^ and 1 · 10^−2^, as shown in Fig.5c. Being well below the standard threshold for p–value significance of 0.05, these results suggest that the correlation range observed in real swarms is wholly incompatible with a random arrangement of mosquitoes.

#### Individual fluctuations vs mutual synchronization

Having established the relevance of swarm spatial arrangement, we now focus on speed fluctuations. We consider two opposing hypotheses, both of which assume speed localization. In the first, mosquitoes’ individual speed distributions are driven by intrinsic and environmental factors, and variations in speed are solely due to statistical fluctuations. In the second, mosquitoes actively modulate their speed to synchronize with their neighbors.

We mimic the first scenario by generating artificial swarms from our data. As in the previous test, we perform a shuffling procedure; this time we shuffle individual speeds, while preserving the specific spatial configuration of the original swarm. At a given instant of time, we associate the position of each mosquito with a speed randomly chosen from its own distribution, see Fig.6 and Methods. We generate 10^4^ artificial realizations for each swarm and compute their correlations. Note that the core of the localization property is the temporal persistence of the speed fluctuations, specifically the persistence of their signs: if a mosquito is faster than the group average at time *t*, it will likely be faster throughout the entire acquisition. Consequently, if two mosquitoes are correlated at time *t*, i.e. *δs*_*i*_(*t*)*δs*_*j*_(*t*) > 0, their fluctuations at two random instants of time will likely retain the same sign. We therefore expect non-trivial correlations in these artificial swarms. The aim of this test is to assess whether individual fluctuations alone can account for the correlations observed in our swarms, without invoking mutual synchronization.

**Fig. 6.**
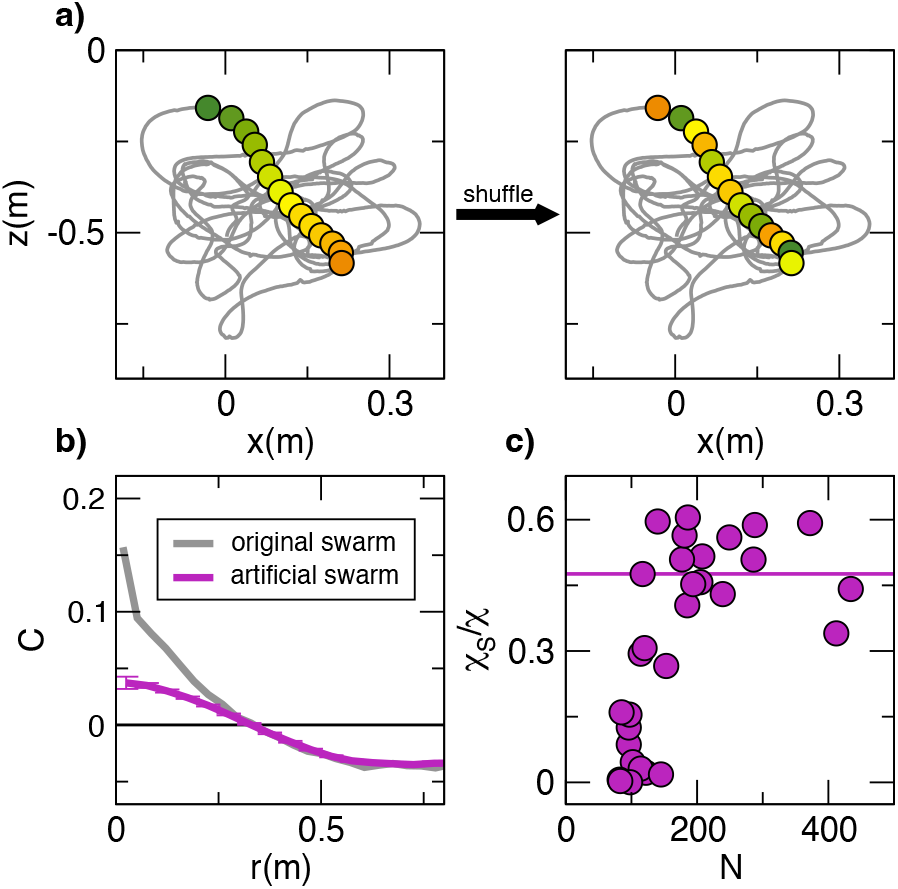
Individual fluctuations vs mutual synchronization. a. A schematic description of the shuffling procedure. For each mosquito in the swarm, we preserve its trajectory (dark gray line), while randomly shuffling its speed values. The dots are a schematic representation of the speed values: in the original swarm, they are ordered in time and colored from green to orange accordingly; after shuffling, speeds are associated to randomly chosen instants of time, as indicated by the mixed colors. **b**. The comparison between correlation in the original (dark gray) and artificial (purple) swarms shows that correlation after shuffling is non-trivial, but cannot account for the full magnitude observed in the of the original data. **c**. The ratio *χ*_*S*_*/χ* between the susceptibility in shuffled and original swarms as a function of the number of mosquitoes in the swarm, *N*. *χ*_*S*_ is significantly lower than *χ*, with the ratio *χ*_*S*_*/χ* on average below 0.5. This ratio reaches a plateau at around 0.5, indicating that a substantial fraction of the correlation in real swarms must then derive from the original temporal dependence of individual speeds, namely from the mutual synchronization of their fluctuations and cannot be explained solely by individual fluctuations.

While correlations in artificial swarms are indeed non-trivial (see Fig.6b), their susceptibility *χ*_*S*_ (see Methods) is significantly lower than that of real swarms *χ*, with the ratio *χ*_*S*_*/χ* on average smaller than 0.5, see Fig.6c. This ratio increases with the size of the swarm, being negligible in small swarms, where the speed localization is weak, and reaching a plateau at around 0.5 for large swarms. A substantial portion of the correlation in real swarms must then derive from the original temporal dependence of individual speeds, namely from the mutual synchronization of their fluctuations.

Altogether, the results presented in this section prove that the correlation in our swarms is not a spurious effect of environmental perturbations or localization; rather, it is the expression of a genuine interaction between individuals, taking the form of speed synchronization based on spatial proximity.

In concluding this Section, let us emphasize that the key to our results lies in the nature of our dataset, which includes a variety of large swarms. As we show in the SI, had our analysis been restricted to small swarms (from 10 to 80 mosquitoes), it would not have been possible to detect the signature of interaction, nor the scale-free property of the correlation.

## 3 Discussion

With this work, we move a step forward in the understanding of swarming behavior in lab-adapted mosquitoes *Anopheles gambiae* (strain G3). Despite the potential deleterious effects of generations of laboratory rearing [20, 39–42], these mosquitoes exhibit robust swarming activity, forming swarms of hundreds of individuals (up to 400 in the data presented here), well within the range observed in field experiments [43, 44].

In contrast to most studies on swarming [9, 11, 15, 16, 18, 20, 21, 45–47], which focus on positional (attraction/repulsion) or directional (alignment) interactions, we provide experimental evidence of male-male interaction in the form of speed synchronization. This positions laboratory mosquito swarms within a framework distinct from previously studied systems, such as bird flocks, fish schools and, most notably, midge (*Diptera Chironomidae*) field swarms [4, 5, 11]. In those systems, correlation is primarily observed in the velocity domain, involving the direction of motion, and it is accompanied by speed correlation (see SI for an example comparing velocity and speed correlation in midge swarms). Conversely, our data exhibit strong correlation in speed, but no significant correlation in velocity (see SI). This discrepancy may stem from biological factors, specifically from interaction mechanisms that depend on the system’s taxonomic class and species. But it is also plausible that environmental conditions — laboratory versus field — influence dominant behavioral traits, such that some aspects of interaction are inhibited under laboratory conditions. A case of interest is that of midges, where velocity correlations are non–trivial in field experiments [11], while not significant in the controlled environment of the laboratory [48]. This could either confirm the discrepancy between laboratory and field behavior or arise from experimental limitations, such as the size of laboratory swarms, which may not be large enough to effectively detect the emergence of collective behavior (see SI).

A natural step for future work is then to benchmark our results against wild-type and field-derived *Anopheles gambiae*, in order to evaluate the extent to which this behavior reflects natural swarming and assess the impact of the laboratory environment. Future research should potentially extend to other major *Anopheles* vectors relevant for malaria transmission, such as *Anopheles coluzzii, Anopheles arabiensis*, and *Anopheles funestus*, for which species-specific swarming behavior has been described [49–52].

The two key features we identify - localization and speed correlation - require the design of novel models for swarm dynamics. Standard models assume an interaction acting on the velocity and homogeneity among individuals [29], such that differences, at a fixed instant in time, are solely due to statistical fluctuations. Localization instead implies heterogeneity and temporal persistence: individuals exhibit characteristic positions and speeds, which are slightly different from those of the other members of the group, and are maintained over time. From a biological perspective, localization may be seen as a natural property of the system. Environmental and genetic factors lead to differentiation of individual traits, which results in specific kinematic properties. As with humans - who may walk at different speeds depending on factors such as age, weight, or overall physical condition — mosquitoes could also exhibit preferences for specific locations and speeds. While these individual preferences may explain the presence of localization, our results demonstrate that they cannot account for the correlation found in our swarms. Consequently, a more sophisticated mechanism, based on mutual adaptation of their speed, must be at play in our swarms.

The lack of speed spatial structure provides a powerful insight into this mechanism. It excludes the possibility that mosquitoes choose their positions to match their intrinsic preferred speeds with those of neighboring individuals. Such a process would imply the formation of speed–based clusters, where fast and slow mosquitoes occupy distinct regions of the swarm, a feature that we do not find in our data. It is therefore more plausible that mosquitoes choose their positions based on environmental cues or, more likely, through positional interactions such as mutual attraction. While we cannot definitively prove the latter from our data, our results suggest that, once settled within the swarm, mosquitoes are able to modulate their speeds to synchronize with their neighbors, regardless of their individual preferences, much as people walking together. Through a mechanism that may be modeled as an imitation process, they modulate their speed to reduce differences with their neighbors, similarly to what is observed in bird flocks and fish schools [4, 5].

This imitation mechanism may appear to contradict speed localization: if mosquitoes synchronize their speed, they should theoretically converge toward the same speed distribution, hence they should not exhibit localization. This apparent contradiction may be explained with an interaction mechanism with low resolution. Due to mechanosensory constraints, mosquitoes may not be able to perceive the difference in speed from that of their neighbors, when it falls below a certain perception threshold. As a result, only when the difference exceeds this threshold, mosquitoes perceive themselves as out of sync and adjust their speed — such that the differences fall below the perception threshold. The system can therefore show synchronization and, at the same time, allow mosquitoes to keep their individuality within the swarm.

An interaction based on the speed, combined with a perception threshold, could serve as a effective mechanism when designing a model. But from a biological perspective, there is an alternative interpretation of our data: speed correlation may not emerge from an interaction directly acting on speed itself, but rather may be an epiphenomenon of a different kind of interaction, which indirectly forces individual speeds to be synchronized.

The parallel with the acoustic mechanism is compelling, but still not robustly supported by experimental data. The mosquito auditory system naturally presents the perception threshold we introduced above to reconcile localization and correlation [23], and a form of acoustic interaction has been already documented in small arrays of tethered *Aedes aegypti* mosquitoes, [27]. More interestingly, mosquitoes play an active role in shaping the sound they perceive [53], in such a way that, by modulating their own wing beat frequencies, they can effectively modulate the environmental sounds to which they can respond. Through an acoustic interaction, males could then transform the noisy environment of the swarm, generated by the sounds of hundreds of individuals, into one where only biologically relevant signals (i.e., those from females) stand out, thereby preserving mating efficiency especially in the most acoustically complex large swarms [44].

The link between acoustic and speed interaction lies in mosquitoes’ wing beat frequency, which defines the sound emitted and, at the same time, influences flight speed. A correlation between wing beat frequency and speed appears reasonable, and consistent with experimental observations in both laboratory [54] and field swarms [55], where simultaneous changes in wing beat frequency and speed are documented at a group level. However, empirical data on the individual speed fluctuations in relation to wing beat frequency modulation are still lacking, as they require sophisticated measures of individually emitted sounds in combination with high-speed video recording. This gap highlights the need for new experiments aimed at determining whether wing beat frequency and speed are positively correlated when mosquitoes participate in a swarm — an essential step to confirm our biological interpretation.

## 4 Methods

In this section, we aim to provide all the formal definitions and computational details related to the results presented in Section 2. To facilitate the reader, the outline of this section reproduces the structure of Section 2, with a first subsection on the experiments, a second on the swarm localization properties, and a third on the experimental evidence of interaction.

### 4.1 Experiments

We performed 3*D* swarming experiments on mosquitoes *Anopheles gambiae* (*An. gambiae*) G3 strain (MRA-112). Mosquitoes were contained in a containment level 2 facility at the Department of Medicine and Surgery, Perugia, Italy, authorization N. PG/IC/Imp2/13/001-Rev2 from Ministry of Health.

As in [20], mosquitoes are reared using the MR4 protocol [56], with temperature at 28 ± 1°C and 75 ± 10% relative humidity under a 12*/*12 – *h* light/dark cycle. Details on feeding procedure and on the husbandry techniques may be found in [20].

Males and females are manually separated as pupae by microscopic examination of the terminalia and raised in separate cages, ensuring all males are virgins. For each experiment, between 800 and 1800 2 − 3-day-old males were released from ‘Bug-Dorm’ type cages approximately 1 h before the onset of the artificial sunset, and left in the large swarming cage containing multiple resting sites and sources of sugar/honey solution as described previously in [20], for the next few days for data collection.

#### Swarming arena setup

As described elsewhere [20], data collection took place in the Insectary Field 1 (measuring 8.5*m* × 3.8*m* × 3*m, L* × *W* × *H*) at the Department of Medicine and Surgery of the University of Perugia, which contains one large cage of 5.0*m* × 3.5*m* × 2.6*m*). Since swarms naturally occur at dusk, the swarming arena is illuminated using six 850nm IR LED lamps (RAYMAX 300). We reproduced the visual stimuli needed for mosquitoes to swarm; specifically, see [20]: i. a contrasting ground marker (55*cm* × 55*cm*), at approximately 1.1m from the back wall; ii. two series of lights: three 2700*K* 8W lights located on the floor outside the cage; iii. three 2700K 9W LED lights placed on the floor at the back of the cage to simulate twilight and an artificial horizon, see [20] for a detailed scheme of the setup.

#### Data collection

We collect 3*D* data with a system of high–speed synchronized cameras (IDT-MotionScope M5) equipped with Schneider Xenoplan 28mm f/2.0 optics. Each camera is placed on a plexiglass shelf, supported by a metal tripod (Manfrotto Pro Digital Tripod 475B), allowing the correct orientation of the cameras towards the swarming area. The three cameras are located outside of the large cage at a distance of 3.5 − 3.9m from the ground marker. Cameras record the same swarm from different positions, so that, using the 3*D* tracking software GReTA [28], we are able to reconstruct three dimensional trajectories of each individual in the swarm. We calibrate the mutual position and orientation of the cameras after each experiment, following the post-calibration procedure defined in [57]; camera internal parameters (focal length, distortion coefficient and image center) are instead calibrated using a standard method [58]. As indicated in the text and in Supp. Table 1, two modes of data acquisition were employed: 1. using a frame rate of 170fps of duration approx. 15.9s 2. using a frame rate of 80fps and duration approx. 33 s.

#### Notations

We summarize here the notations we will use in the following two subsections:

- *N* represents the number of mosquitoes in the swarm;
- *i* = 1, …, *N* refers to a specific individual in the swarm;
- *T*_*i*_, with *i* = 1, …, *N*, refers to the duration of the trajectories of mosquito *i*, which is not always as long as the entire acquisition (*T*_*i*_ ≤ *T*);
- 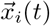 is the 3*D* position vector of mosquito *i* at time *t*, expressed in m. In terms of the 3*D* coordinates, 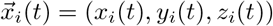;
- 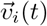 is the 3*D* velocity vector of mosquito *i* at time *t*, expressed in m/s, and computed from the spatial displacement between two consecutive frames with the following expression:

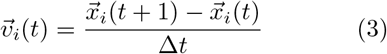

with Δ*t* = 1*/*80s or Δ*t* = 1*/*170s, depending on the data;
- *s*_*i*_(*t*) is the speed of mosquito *i* at time *t*, namely the magnitude velocity vector *s*_*i*_(*t*) = |*v*_*i*_(*t*)|.

### 4.2 Localization within the swarm

#### Spatial localization

We quantify the spatial localization in our swarms, by computing group and individual spatial distributions on the horizontal plane and along the vertical axis.

#### Group spatial distributions - Swarm extent

We compute the swarm centroid, 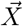, as the average over the position of all mosquitoes at all instants of time:

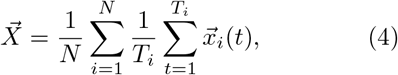

and 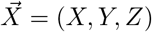

We measure the distance of each mosquito from 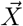 on the horizontal plane:

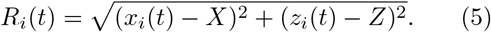

On the set of all these distances, we define the group probability distribution, *P*_ℛ_ (*R*), the one highlighted in dark gray in Fig.1c.

We measure the radius of the swarm as the mean value, *R*, of the distribution *P*_ℛ_ (*R*):

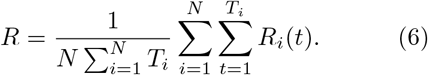

*R* varies from swarm to swarm, ranging from 0.19m to 0.45m, see Table S1.

We measure the distance of each mosquito from 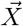 along the *y*-axis:

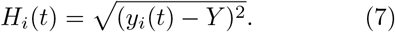

On the set of all these distances, we define the group probability distribution, *P*_ℋ_ (*H*), the one highlighted in dark gray in Fig.1h.

We measure the elongation of the swarm as the mean value, *H*, of the distribution *P*_ℋ_ (*H*):

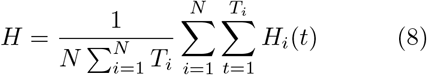

*H* varies from swarm to swarm, ranging from 0.10m to 0.19m, see Table S1.

#### Individual spatial distributions - Trajectory extent

We compute the centroid of each trajectory, 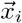, by averaging the position of each individual over time:

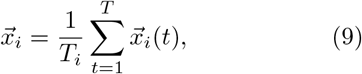

and 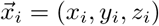.

We then compute the distances of each mosquito from its own centroid, which we denote by *r*_*i*_(*t*):

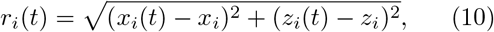

when the distance is computed on the horizontal plane, and by *h*_*i*_(*t*):

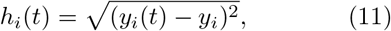

when the distance is computed on the vertical axis.

We define the radius of trajectory *i* as:

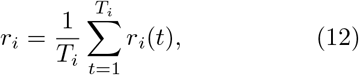

and the elongation of trajectory *i* as:

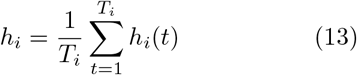

In Fig.1d,i we show, with dark purple circles, the set of *r*_*i*_ and *h*_*i*_ for all the mosquitoes in one of our swarms. With the light purple shading we are instead showing their standard deviation:

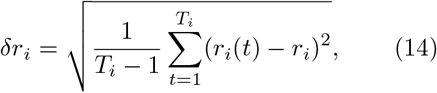

and

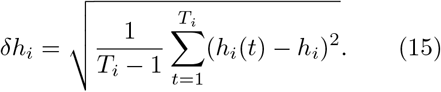

Finally, we aggregate all individual distances *r*_*i*_(*t*) and *h*_*i*_(*t*) obtaining the two individual distributions, *P*_*r*_(*r*) and *P*_*h*_(*h*), shown in purple in Fig.1c,h. For each swarm, we then define the typical trajectory radius, *r*, and the typical elongation, *h*, as:

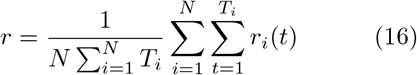

and

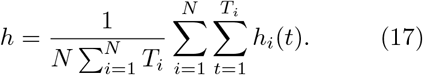

We compare the spatial extent of individual trajectories with that of the swarm, by computing the two ratios *r/R* (from eq.(16) and eq.(6)) and *h/H* (from eq.(17) and eq.(8)) that we show in Fig.1e,j as a function of the number of mosquitoes in the swarm.

#### Speed localization

We consider the set 𝒮_*S*_ = {*s*_*i*_(*t*), *i* = 1, …, *N* and *t* = 1, …, *T}*, of the speeds of all mosquitoes at all instants of time. From this set, we define the group probability distribution, *P*_*S*_(*s*), (highlighted in dark gray in Fig.2a).

With a similar procedure we can also define individual probability distributions. We consider the sets 𝒮_*i*_ = {*s*_*i*_(*t*), *t* = 1, …, *T}*, of the speed of mosquito *i* at all instants of time. From these sets, we define individual probability distributions, 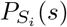, one for each individual. In Fig.2a, we show the distributions of three different mosquitoes in purple.

For each mosquito *i*, we compute the mean speed as:

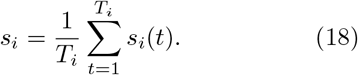

We define the distribution of these mean values, which we show in Fig.2b, where in light purple we highlight its standard deviation *σ*_*S*_.

We also estimate the standard deviation of the mean from the group distribution, as the standard error, Σ:

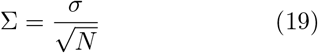

where with *σ* we define the standard deviation of the group speed distribution. In the absence of speed localization, the two values (*σ*_*S*_ and Σ) should be compatible, and variations in individual mean speeds would reflect only statistical fluctuations. This is because in the absence of localization, individual and group speed localization follow the same distribution. Therefore, the standard deviation of each mosquito is equal to *σ*. Since 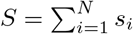.

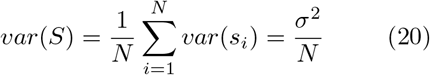

where *var*(*S*) and *var*(*s*_*i*_) denotes the variance of the mean and the variance of individual speeds respectively.

#### Probability of positive speed fluctuations

To better quantify the localization, we move from absolute speed to their fluctuations with respect to the swarm average. To this aim, at each instant in time, we define the swarm average as:

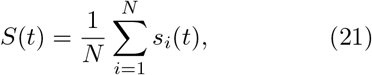

and the individual speed fluctuations for mosquito *i* as:

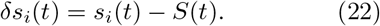

From the set of all the fluctuations, we compute the probability distribution, see inset of Fig.2d. By definition the mean value of this distribution is equal to 0m/s.

For each individual, we compute the mean of its fluctuation, *δs*_*i*_, as:

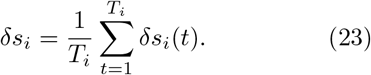

Surprisingly, these individual mean values are generally different from 0m/s, and are not symmetric with respect to the zero.

We give a measure of the individual asymmetry by computing, for each mosquito, the proportion of time that it spends having positive fluctuations. These quantity, denoted by *P*_0*i*_, is computed as follows:

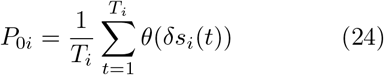

where *θ*(*δs*_*i*_) is the Heaviside step function defined as:

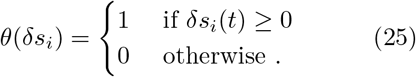

In the absence of localization, *P*_0*i*_ should be equal to 0.5: a mosquito should spend half of the time flying faster than the group average (positive fluctuations) and half of the time flying slower (negative fluctuations). We find instead that in our swarms, it spans the entire range between 0.1 and 0.9, as shown in Fig.2d, where we plot the values of *P*_0*i*_ as a function of *δs*_*i*_.

### 4.3 Experimental evidence of male–male interaction

#### Spatial correlation functions of the speed fluctuations

We start by computing the speed fluctuation of each individual at each instant in time, as in eq.(22). Then, for each pair of mosquitoes (*i, j*), we compute:

- the correlation coefficient defined as in eq.(2), *c*_*ij*_(*t*) = *δs*_*i*_(*t*) · *δs*_*j*_(*t*);
- the mutual distance 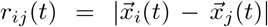 where | · | indicates the vector magnitude.

We then define the correlation at time *t* as:

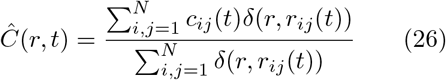

where *δ*(*r, r*_*ij*_) is the Kronecker delta function:

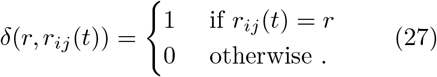

With the definition in eq(26), we are averaging the correlation coefficients over pairs at mutual distance equal to *r*. Specifically, the factor *δ*(*r, r*_*ij*_(*t*)) selects only those pairs for which *r*_*ij*_(*t*) = *r*, neglecting the contribution of all other pairs. In the numerator, we are then summing the correlation coefficient of all those pairs, while in the denominator we are counting how many they are.

The *Ĉ* (0, *t*) is a special case: at a fixed instant in time, the only pairs that participate in *Ĉ* (0, *t*) are the ones at a mutual distance *r* = 0, hence only the pairs where *i* = *j*:

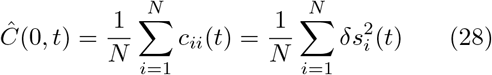

which is the average square fluctuation of the speed at time *t*.

We use *Ĉ* (0, *t*) to define the normalized correlation function at time *t, C*(*r, t*), as follows:

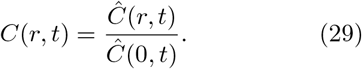

This normalized correlation function is such that *C*(0, *t*) = 1.

Finally, we average over time and obtain the spatial correlation function we show in Fig.3:

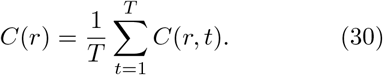

When working on finite systems, the definition of the correlation function in eqs.(26-29) has two limitations. The first is that we cannot compute the *C*(*r*) on continuous values: the mutual distances *r*_*ij*_ define a finite set of values (precisely *N* · (*N* − 1)). The second is that if we strictly use the definition in eq.(26), we will not have statistically relevant samples, because for each value of *r* we will have essentially only one pair: it is very unlikely that two (or more) pairs are located exactly at the same mutual distance.

To overcome these two issues, we perform a binning in *r* of the *Ĉ* (*r, t*), and consequently of the *C*(*r*). More specifically, we fix the size of the bin, Δ*r*, and we define the discrete correlation function as:

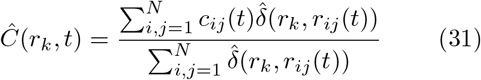

where:

- *k* = 1, …, *k*_*M*_ ;
- *k*_*M*_ is defined as the smallest integer such that *k*_*M*_ Δ*r* is greater or equal to the size of the swarm, *L*, hence *k*_*M* − 1_Δ*r* < *L* ≤ *k*_*M*_ Δ*r*;
- 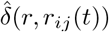 is a Kronecker delta function modified as follows:

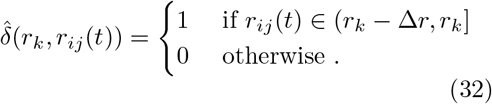

For *r* = 0, we define the discrete correlation function as the continuous one defined in eq.(28).

We finally average over time, and obtain:

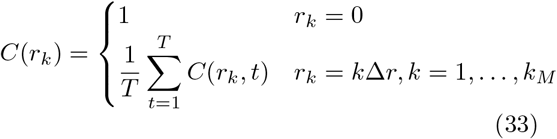

We are essentially keeping the correlation for *r* = 0 separated from the other bins, since it has a special meaning. In the other bins we are instead averaging the correlation coefficient over intervals of size Δ*r*. The choice of Δ*r* is relevant here: if it is too small, the correlation function would be dominated by noise (as occurs in the continuous case); if it is too large, we would not be able to detect a correlation signal. Our choice is to define Δ*r* as half of the typical nearest neighbor distance in the system.

#### Nearest neighbor distance, d_NN_

We compute the distance of each mosquito from the other *N* − 1, and we select the smallest. We do this, for all the mosquitoes in the swarm and for all the instants of times. We define the typical distance to the nearest neighbor, *d*_*NN*_, as the median value of this set.

#### Correlation range, r_0_

We compute the correlation range as the first point where the correlation function crosses the *x*-axis. To this aim, we find the smallest *k* such that *C*(*r*_*k*_) > 0 and *C*(*r*_*k*+1_) < 0, and we estimate *r*_0_ with a linear interpolation between the two points (*r*_*k*_, *C*(*r*_*k*_)) and (*r*_*k*+1_, *C*(*r*_*k*+1_)).

#### Correlation length, *ξ*

Following [4] we computed the correlation length using the following integral definition:

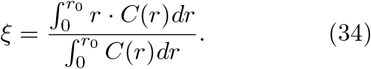

where *r*_0_ is the correlation range. This definition has two main benefits: the first is that it uses not only the point where the correlation function crosses the *x*-axis, but the entire shape of the correlation function (up to *r*_0_); the second, which is also the most relevant, is that this definition is general and it may be applied to correlation functions in different regimes. We compute the two integrals in eq.(34) with the trapezoid rule.

#### Swarm size, L

For consistency with [4, 11] we compute the size of the swarm *L* as the maximum distance between two mosquitoes. Specifically, at each instant in time we measure the distances between each pair of mosquitoes and we define the size at that frame, *L*(*t*), as the maximum distance. We finally define *L* as the median value of the set of all these *L*(*t*).

#### Standard deviation for *ξ* and L

We measure the standard deviation on the correlation length *ξ* and on the size of the swarm *L*, with a resampling method, see [59], already used in [18]. We randomly select half of the *T* different frames of the acquisition and compute, on this sub-sample, the correlation function, the correlation length and the size of the swarm. We repeat this procedure 10^5^ times, obtaining a set of values for *ξ* and *L*. We extracted the standard deviations of the two sets, which are the estimates of the standard deviation for *ξ* and *L* shown in Fig.3d.

#### Evidence of male–male interaction

In this section we provide the details of the different techniques we used to prove that the observed correlation is a genuine signature of interaction in our swarms.

#### Lack of speed spatial structure

To detect speed based clusters, we apply a connected components technique, where two mosquitoes are assigned to the same cluster if the difference between their speed fluctuations is below a specified threshold. This threshold is determined independently for each swarm using a standard *elbow* method. Specifically, for each mosquito *i* at time *t*, we measure the difference with all the other members of the swarm:

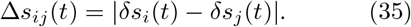

with *δs*_*i*_(*t*) and *δs*_*j*_(*t*) denote the speed fluctuation at time *t* of mosquito *i* and *j*.

For each mosquito *i*, we then pick the minimum difference Δ*s*_*i*_ = *min*_*i*!*neqj*_Δ*s*_*ij*_(*t*). We pool the minimum distances across all time and all mosquitoes. Finally we sort these values in ascending order, obtaining a curve that exhibits a characteristic *elbow*. The threshold is set at this elbow, providing a threshold specific to each swarm, typically equal to 0.051m/s - in the range between 0.028m/s and 0.076m/s.

For each frame, we identify the primary cluster, as the largest one. On the secondary clusters we run a second clustering procedure, this time, in space. We apply a connected components clustering technique, again choosing the threshold with the elbow method and fixing the minimum number of points in each cluster equal to 4, following a standard approach for three dimensional data. The spatial elbow is typically equal to 0.25m - in the range between 0.17m and 0.28m.

#### Susceptibility

To quantify the correlation in the system, we use the susceptibility *χ* defined as:

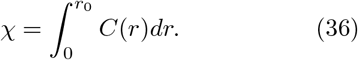

This variable integrates the correlation function up to *r*_0_, and it is larger the higher the correlation, thereby giving a measure of the amount of correlation in the system.

#### Relevance of spatial arrangement

We used a shuffling procedure to generate non–interacting and realistic swarms, directly from the data. These artificial swarms present the same localization properties as real swarms, from which they differ only in the spatial arrangement of mosquitoes.

In eq.(9), we defined individual centroids as the average over time of the position of single mosquitoes. We can now rewrite the position at time *t* of a mosquito *i*, as a function of its centroid:

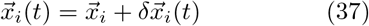

where 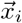 is the trajectory centroid, hence not– dependent on time, and 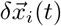 represents the location of mosquito *i* at time *t* with respect to its own centroid.

We have then a set of centroids, 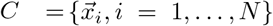, each associated with a specific mosquito. In the shuffling procedure, we scramble the associations between centroids and mosquitoes. For instance, we associate the centroid of mosquito *j* to mosquito *i*, and we define a new trajectory,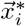 such that:

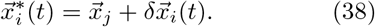

In doing so, the position of the new trajectory is shifted in space, in such a way that its new centroid is 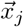 instead of 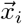. But, its velocity and its speed in time are not affected by this shift, as well as its individual spatial distribution.

We repeat this procedure for all the mosquitoes in the swarm, associating each centroid with a different trajectory. In the example of Fig.5, the yellow trajectory is shifted on the turquoise centroid, the purple trajectory on the yellow centroid, and the turquoise trajectory on the yellow centroid.

At the end of the shuffling process, we have an artificial swarm where, by mixing up mosquito positions, we cut potential interaction links due to spatial proximity. But at the same time, we are not modifying individual spatial extent nor speed, which will both present the same localization properties as the original real swarm.

We also compute the probability of finding *r*_0*S*_ greater or equal to *r*_0_. To this aim, we measured the ratio between the occurrences of the shuffled swarms with *r*_0*S*_ ≥ *r*_0_ and the total number of times we shuffled the swarm (10^6^). This probability is essentially the p–value for *r*_0_.

#### Relevance of equal-time speed fluctuations

Similarly to the previous test, we used a shuffling procedure to generate non–interacting and realistic swarms, directly from the data. This time, artificial swarms present the same spatial arrangement of real swarms, from which they differ only in the temporal dependence of individual speeds.

Specifically, we randomly shuffle over time the temporal sequence of the speed of each individual mosquito. As a result, the position of mosquito *i* at time 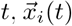, which was originally associated to the speed *s*_*i*_(*t*), is instead paired with the speed of the same individual at a different, randomly chosen instant *t*^*′*^, *s*_*i*_(*t*^*′*^). This procedure decouples individual positions from their instantaneous speeds and, more importantly, disrupts the simultaneity of speed fluctuations across the group: while 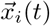 is associated to *s*_*i*_(*t*), the position of mosquito *j* at the same instant of time *t* is associated to its speed at time *t*^*′′*^.

We repeat this procedure 10^4^ times for each swarm. At each run, we compute correlation functions that are finally averaged. On the resulting average correlation we compute the susceptibility, *χ*_*S*_, that we compare with the correlation of the original swarm, *χ*.

## Author contributions

SM, LP, MP, RS conceived and designed the experiments.

GI, ML performed the experiments. SM analyzed the data.

MF, FG, AL, ML, LP, GZ contributed material/-analysis tools.

SM wrote the paper.

## Acknowledgments

We thank Andrea Cavagna and Irene Giardina for discussions and for financial support, both in terms of resources and equipment. We thank CoBBS group, to which MF, FG, SM and LP belong, for the data on the midge swarm presented in SI. SM thanks Federica Ferretti, who unwittingly triggered part of this work with a question at the IntCha2025 conference. This work was supported by ERC Grant RG.BIO (Contract n. 785932), and MIUR Grant PRIN2022 funded by the European Union – Next Generation EU, under Italy’s National Recovery and Resilience Plan (PNRR), Mission 4, Component 1 (CUP B53D23011780006).

## Data and code availability

Requests for further information and resources should be directed to and will be fulfilled by the lead contact, Stefania Melillo.

- All data required to evaluate the conclusions of this paper will be made available by the lead contact upon reasonable request;
- All original code developed for this paper will be provided by the lead contact upon request;
- Any additional information required to reanalyze the data reported in this paper is available from the lead contact upon request.

## References

[1] Sarfati, R., Hayes, J.C., Peleg, O.: Self-organization in natural swarms of Photinus carolinus synchronous fireflies. Science Advances 7(28) (2021)

[2] Strogatz, S.H., Stewart, I.: Coupled oscillators and biological synchronization. Scientific american 269(6), 102–109 (1993)

[3] Attanasi, A., Cavagna, A., Del Castello, L., Giardina, I., Grigera, T.S., Melillo, S., Parisi, L., Pohl, O., Shen, E., Viale, M.: Information transfer and behavioural inertia in starling flocks. Nature Physics 10 (2014)

[4] Cavagna, A., Culla, A., Feng, X., Giardina, I., Grigera, T.S., Kion-Crosby, W., Melillo, S., Pisegna, G., Postiglione, L., Villegas, P.: Marginal speed confinement resolves the conflict between correlation and control in collective behaviour. Nature Communications 13 (2022)

[5] Katz, Y., Tunstrøm, K., Ioannou, C.C., Huepe, C., Couzin, I.D.: Inferring the structure and dynamics of interactions in schooling fish. Proceedings of the National Academy of Sciences 108(46), 18720–18725 (2011)

[6] Herbert-Read, J.E., Perna, A., Mann, R.P., Schaerf, T.M., Sumpter, D.J.T., Ward, A.J.W.: Inferring the rules of interaction of shoaling fish. Proceedings of the National Academy of Sciences 108(46), 18726–18731 (2011)

[7] Sumpter, D.J.T.: Collective Animal Behavior. Princeton University Press, Princeton (2011)

[8] Strogatz, S.: Sync: The Emerging Science of Spontaneous Order. Hyperion, New York (2003)

[9] Kelley, D., Ouellette, N.: Emergent dynamics of laboratory insect swarms. Scientific reports 3, 1073 (2013)

[10] Gorbonos, D., Puckett, J., Vaart, K., Sinhuber, M., Ouellette, N., Gov, N.: Pair formation in insect swarms driven by adaptive long-range interactions. Journal of the Royal Society, Interface 17 (2020)

[11] Attanasi, A., Cavagna, A., Del Castello, L., Giardina, I., Melillo, S., Parisi, L., Pohl, O., Rossaro, B., Shen, E., Silvestri, E., Viale, M.: Collective behaviour without collective order in wild swarms of midges. PLoS Comput Biol 10(7) (2014)

[12] Downes, J.A.: The swarming and mating flight of diptera. Annual Review of Entomology 14, 271–298 (1969)

[13] Charlwood, J.D., Jones, M.D.R.: Mating behaviour in the mosquito, anopheles gambiae s.1.save. Physiological Entomology 4(2), 111–120 (1979)

[14] Baeshen, R.: Swarming behavior in anopheles gambiae (sensu lato): Current knowledge and future outlook. Journal of Medical Entomology 59(1), 56–66 (2021)

[15] Cribellier, A., Poda, B.S., Diabaté, A., Roux, O., Muijres, F.T.: The complex swarming dynamics of malaria mosquitoes emerges from simple minimally-interactive behavioral rules. bioRxiv (2024)

[16] Attanasi, A., Cavagna, A., Del Castello, L., Giardina, I., Melillo, S., Parisi, L., Pohl, O., Rossaro, B., Shen, E., Silvestri, E., Viale, M.: Finite-size scaling as a way to probe near-criticality in natural swarms. Phys. Rev. Lett. 113 (2014)

[17] Cavagna, A., Conti, D., Creato, C., Del Castello, L., Giardina, I., Grigera, T.S., Melillo, S., Parisi, L., Viale, M.: Dynamic scaling in natural swarms. Nature Physics 13 (2017)

[18] Cavagna, A., Di Carlo, L., Giardina, I., Grigera, T.S., Melillo, S., Parisi, L., Pisegna, G., Scandolo, M.: Natural swarms in 3.99 dimensions. Nature Physics 19 (2023)

[19] Manoukis, N., Diabate, A., Abdoulaye, A., Diallo, M., Dao, A., Yaro, A., Ribeiro, J., Lehmann, T.: Structure and dynamics of male swarms of anopheles gambiae. Journal of medical entomology 46, 227–35 (2009)

[20] Cavagna, A., Giardina, I., Gucciardino, M.A., Iacomelli, G., Lombardi, M., Melillo, S., Monacchia, G., Parisi, L., Peirce, M.J., Spaccapelo, R.: Characterization of lab-based swarms of anopheles gambiae mosquitoes using 3d-video tracking. Scientific Reports 13 (2023)

[21] Poda, B.S., Cribellier, A., Feugére, L., Fatou, M., Nignan, C., Sales Hien, D.F., Müller, P., Gnankiné, O., Dabiré, R.K., Diabaté, A., Muijres, F.T., Roux, O.: Spatial and temporal characteristics of laboratory-induced Anopheles coluzzii swarms: Shape, structure, and flight kinematics. iScience 27(11) (2024)

[22] Gibson, G., Russell, I.: Flying in tune: Sexual recognition in mosquitoes. Current Biology 16(13), 1311–1316 (2006)

[23] Somers, J., Georgiades, M., Su, M.P., Bagi, J., Andrés, M., Alampounti, A., Mills, G., Ntabaliba, W., Moore, S.J., Spaccapelo, R., Albert, J.T.: Hitting the right note at the right time: Circadian control of audibility in anopheles mosquito mating swarms is mediated by flight tones. Science Advances 8(2) (2022)

[24] Poda, B., Nignan, C., Gnankine, O., Dabiré, R., Diabate, A., Roux, O.: Sex aggregation and species segregation cues in swarming mosquitoes: Role of ground visual markers. Parasites & Vectors 12 (2019)

[25] Butail, S., Manoukis, N., Diallo, M., Ribeiro, J., Paley, D.: The dance of male Anopheles gambiae in wild mating swarms. In: Journal of Medical Entomology (2013)

[26] Shishika, D., Manoukis, N., Butail, S., Paley, D.: Male motion coordination in anopheline mating swarms. Scientific reports 4 (2014)

[27] Aldersley, A., Champneys, A., Homer, M., Bode, N.W.F., Robert, D.: Emergent acoustic order in arrays of mosquitoes. Current Biology 27(22), 1208–1210 (2017)

[28] Attanasi, A., Cavagna, A., Del Castello, L., Giardina, I., Jelic, A., Melillo, S., Parisi, L., Pellacini, F., Shen, E., Silvestri, E., et al.: Greta -a novel global and recursive tracking algorithm in three dimensions. Pattern Analysis and Machine Intelligence, IEEE Transactions on 99 (2015)

[29] Shaebani, M.R., Wysocki, A., Winkler, R.G., Gompper, G., Rieger, H.: Computational models for active matter. Nature Reviews Physics 4 (2020)

[30] Clements, A.N.: The Biology of Mosquitoes: Volume 2: Sensory Reception and Behaviour. CABI Publishing, Wallingford (1999)

[31] Yuval, B.: Mating systems of blood-feeding flies. Annual Review of Entomology 51(1), 413–440 (2006)

[32] Fawaz, E.Y., Allan, S.A., Bernier, U.R., Obenauer, P.J., Diclaro, J.W.: Swarming mechanisms in the yellow fever mosquito: aggregation pheromones are involved in the mating behavior of aedes aegypti. Journal of Vector Ecology 39(2), 347–354 (2014)

[33] South, A., Catteruccia, F.: Sexual selection and the evolution of mating systems in mosquitoes. In: Advances in Insect Physiology vol. 51, pp. 67–92. Elsevier, London (2016)

[34] Göpfert, M.C., Robert, D.: Nanometre-range acoustic sensitivity in male and female mosquitoes. Proc Biol Sci. 267 (2000)

[35] Nakata, T., Simões, P., Walker, S.M., Russell, I.J., Bomphrey, R.J.: Auditory sensory range of male mosquitoes for the detection of female flight sound. Journal of The Royal Society Interface 19(193), 20220285 (2022)

[36] Gibson, G., Warren, B., Russell, I.: Humming in tune: Sex and species recognition by mosquitoes on the wing. Journal of the Association for Research in Otolaryngology 11 (2010)

[37] Cavagna, A., Giardina, I., Grigera, T.S.: The physics of flocking: Correlation as a compass from experiments to theory. Physics Reports 728, 1–62 (2018). The physics of flocking: Correlation as a compass from experiments to theory

[38] Reynolds, A.M.: Phase transitions in insect swarms. Phys Biol. 20(5) (2023)

[39] Catteruccia, F., Masiga, D., Talesa, V., Mireji, P., Paton, D., Thailayil, J., Achinko, D.: Swarming and mating activity of Anopheles gambiae mosquitoes in semi-field enclosures. Medical & Veterinary Entomology 30(1), 14–20 (2016)

[40] Paton, D., Touré, M., Sacko, A., Coulibaly, M.B., Traoré, S.F., Tripet, F.: Genetic and environmental factors associated with laboratory rearing affect survival and assortative mating but not overall mating success in Anopheles gambiae sensu stricto. PLoS One 8(12) (2013)

[41] Nignan, C., Poda, B.S., Sawadogo, S.P., et al.: Local adaptation and colonization are potential factors affecting sexual competitiveness and mating choice in Anopheles coluzzii populations. Sci Rep 12(636) (2022)

[42] Facchinelli, L., Valerio, L., Lees, R.S., et al.: Stimulating Anopheles gambiae swarms in the laboratory: application for behavioural and fitness studies. Malar J 14(271) (2015)

[43] Diabaté, A., Yaro, A.S., Dao, A., et al.: Spatial distribution and male mating success of Anopheles gambiae swarms. BMC Evol Biol 11(184) (2011)

[44] Ouédraogo, T.F.X., Sawadogo, S.P., Millogo, A.A., et al.: Characterization of factors influencing swarm dynamics and mating efficiency in Anopheles coluzzii. Parasites Vectors 18(512) (2025)

[45] Ni, R., Ouellette, N.T.: On the tensile strength of insect swarms. Phys. Biol. 13(4) (2016)

[46] Vaart, K., Sinhuber, M., Reynolds, A.M., Ouellette, N.T.: Mechanical spectroscopy of insect swarms. Science Advances 5(7) (2019)

[47] Sinhuber, M., Vaart, K.V., Feng, Y., et al.: An equation of state for insect swarms. Sci Rep 11(3773) (2021)

[48] Vaart, K., Sinhuber, M., Reynolds, A.M., Ouellette, N.T.: Environmental perturbations induce correlations in midge swarms. Journal of The Royal Society Interface 17(164), 20200018 (2020)

[49] Kaindoa, E.W., Ngowo, H.S., Limwagu, A., Mkandawile, G., Kihonda, J., Masalu, J.P., Bwanary, H., Diabaté, A., Okumu, F.O.: New evidence of mating swarms of the malaria vector, Anopheles arabiensis in tanzania. Wellcome Open Res. 2, 88 (2017)

[50] Kaindoa, E.W., Ngowo, H.S., Limwagu, A.J., Tchouakui, M., Hape, E., Abbasi, S., Kihonda, J., Mmbando, A.S., Njalambaha, R.M., Mkandawile, G., Bwanary, H., Coetzee, M., Okumu, F.O.: Swarms of the malaria vector Anopheles funestus in tanzania. Malar J. 18, 1 (2019)

[51] Ng’habi, K.R., Mwasheshi, D., Knols, B.G., et al.: Establishment of a self-propagating population of the african malaria vector Anopheles arabiensis under semi-field conditions. Malar J. 9, 356 (2010)

[52] Niang, A., Maïga, H., Sawadogo, S.P., Konaté, L., Faye, O., Lee, Y., Dabiré, R.K., Diabaté, A., Tripet, F.: Perfect association between spatial swarm segregation and the xchromosome speciation island in hybridizing Anopheles coluzzii and Anopheles gambiae populations. Sci Rep. 12, 1 (2022)

[53] Alampounti, A.C., Georgiades, M., Faber, J., Bagi, J., Andrés, M., Bozovic, D., Albert, J.T.: Distortion rules: audibility creation in the mosquito ear. bioRxiv (2025)

[54] Feugére, L., Roux, O., Gibson, G.: Behavioural analysis of swarming mosquitoes reveals high hearing sensitivity in Anopheles coluzzii. J Exp Biol. 5(225) (2022)

[55] Lupshin, D.N., Vorontsov, D.D.: Frequency tuning of swarming male mosquitoes (Aedes communis, Culicidae) and its neural mechanisms. J Insect Physiol. 132, 104233 (2021)

[56] Benedict, M.: Methods in Anopheles Research. MR4 (Malaria Research & Reference Reagent Resource Center), Atlanta, GA, USA (2010)

[57] Cavagna, A., Creato, C., Del Castello, L., Giardina, I., Melillo, S., Parisi, L., Viale, M.: Error control in the set-up of stereo camera systems for 3d animal tracking. The European Physical Journal Special Topics 224(17-18), 3211–3232 (2015)

[58] Zhang, Z.: A flexible new technique for camera calibration. IEEE Transactions on Pattern Analysis and Machine Intelligence 22(11), 1330–1334 (2000)

[59] Leuzzi, L., Marinari, E., Parisi, G.: Probability Theory for Quantitative Scientists. Cambridge University Press, Cambridge (2025)

