## Supplemental Information for "Experimental evidence of male-male interaction in laboratory swarms of *Anopheles gambiae* mosquitoes"

### Supplementary Information for Experimental evidence of male-male interaction in laboratory swarms of *Anopheles gambiae* mosquitoes

Gloria Iacomelli[1◇], Max Lombardi[1,2◇], Leonardo Parisi[2,3◇], Matteo Fiorini[3], Francesca Grucci[3], Alessio Lavorgna[1], Melania Ligato[1], Gaetano Zarcone[1], Matthew J. Peirce[1], Stefania Melillo[2,3†*], Roberta Spaccapelo[1†]

[1]Department of Medicine and Surgery, University of Perugia, Perugia, Italy.
[2]Institute for Complex Systems, National Research Council, Rome, Italy.
[3]Physics Department, Sapienza University, Rome, Italy.

#### 1 Relevance of swarm size in behavioral analysis

In this section, we address a critical issue concerning the relevance of the swarm size, in terms of the number of individuals participating, for the study of collective behavior. In 1972, in one of his seminal works [1], P.W. Anderson stated "the behavior of large and complex aggregates of elementary particles, it turns out, is not to be understood in terms of a simple extrapolation of the properties of a few particles. Instead, at each level of complexity entirely new properties appear". The whole is not greater than the sum of its parts, it is different.

Here, we show that mosquitoes are no exception to this general principle. To this end, we analyze a dataset, distinct from the one we used in the main text, comprising 41 *small* swarms of tens of mosquitoes, ranging from 10 and 80. On this dataset we perform exactly the same analysis presented in the main text, and we ask: if we only had access to this dataset, would we be able to infer the results we presented in the main text from the large swarms dataset? Would we be able to observe correlation and assess its scale–free nature?

Collecting reliable data on large swarms under controlled laboratory conditions requires considerable experimental effort. In our experimental setup, the observed swarm participation ratio was approximately 15% [2], indicating that only a small fraction of the released adult males actively engaged in swarming behavior. Consequently, to achieve a swarm composed of approximately 200–400 active individuals, it is necessary to release at least 3,000 males. Given an expected sex ratio close to 1:1 and accounting for natural mortality during development, this requires rearing approximately 6,000 larvae per experimental replicate. This approach ensures that a sufficient number of viable males can be obtained and manually separated from females at the pupal stage prior to release. Moreover, large-scale swarming experiments require a dedicated spacious cage, which involves both physical space and financial resources.

Being able to infer behavioral rules from small swarms, within a small environment, would help significantly in streamlining the experimental process and reducing costs. Unfortunately, we find this is not the case. As we will discuss in this section, the comparison between small and large

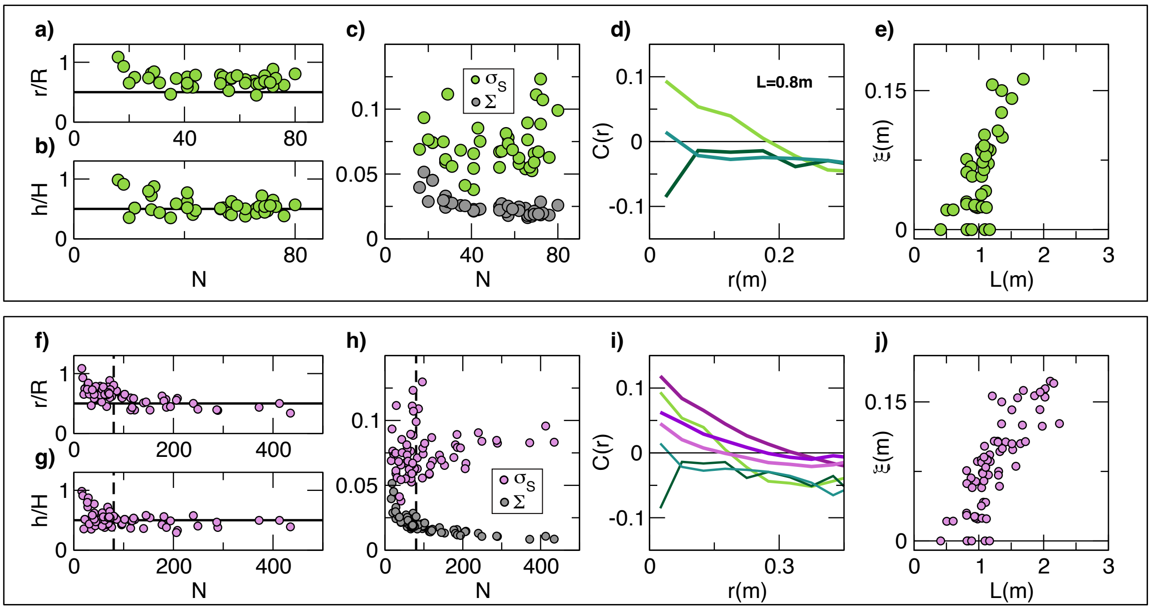


**Fig. S1 Relevance of swarm size in behavioral analysis. a-e)** Localization and correlation analysis on 41 small swarms, in the range between 10 and 80 mosquitoes. **a-b)** The ratio between individual and group radius ($r/R$) and elongation ($h/H$), computed from the individual and group spatial distributions, is larger then what found for large swarm, whose typical value equal to 0.5 is indicated by the horizontal black lines in the two panels. Both $r/R$ and $h/H$ are close to 1 for very small swarms, making spatial localization not evident. **c)** The comparison between the standard error of the speed distribution, $\Sigma$, and the standard deviation of the mean $\sigma_S$, which represents the first hint of speed localization, are not well–separated in small swarms. **d)** Spatial correlation functions of the speed fluctuations for three different swarms of roughly the same size ($L = 0.8$). The three examples does not show a characteristic trend as a function of $r$. Correlation for small distances may be positive or negative, depending on the swarm. **e)** The correlation length, $\xi$, as a function of the size of the system, $L$, does not show the linear dependence we found in large swarms (Pearson coefficient drops to $\rho = 0.61$). In 17 out of 41 swarms (41% of the cases), $\xi$ is equal to 0, meaning that at the typical distance to the nearest neighbor, the correlation is negative. **f-j)** Localization and correlation analysis on 71 swarms: the 30 large swarms presented in the main text and 41 additional small swarms presented in panels a-e. **f-g)** The ratio between individual and group radius ($r/R$) and elongation ($h/H$), computed from the individual and group spatial distributions, decrease with $N$ and reach the plateau equal to 0.5 at approximately $N = 100$. **h)** The comparison between the standard error of the speed distribution, $\Sigma$, and the standard deviation of the mean $\sigma_S$ for the 71 swarms of the dataset shows a clear separation only for large $N$. **i)** The three spatial correlation functions of the speed fluctuations, presented in panel d, are compared with three examples in large swarms. The clear trend observed in large swarms, where the correlation is positive for small distances to smoothly decrease and cross the $x$-axis, is instead not present in small swarms. **j)** The correlation length, $\xi$, as a function of the size of the system, $L$. Including the large swarms presented in the main text, we recover the linear dependence suggesting the scale–free nature of the correlation. Pearson's coefficient $\rho$ is 0.82. only slightly lower than the one obtained considering only large swarms, where it is equal to 0.85. This confirms that small swarms are not adding relevant information to our analysis.

swarms clearly indicates that small swarms are not suitable for the study of collective behavior. Also, if we consider the full dataset, comprising both small and large swarms, the small ones fail to add relevant quantitative and qualitative information. In Fig.S1, we summarize the results we obtain using small swarms in terms of spatial and speed localization, and of the analysis on the speed correlation. The top panel shows results from the small swarms alone, while the bottom panel presents results from the full set of 71 swarms (30 large swarms from the main text and the additional 41 small ones).

**Spatial localization.** In Fig.S1a,b, we show the two ratios $r/R$ and $h/H$, defined in the main text, which measure the ratio between the spatial extent of individual trajectories and of the group, on the horizontal plane and along the vertical axis respectively. Unlike in the large swarm datasets, where both ratios are close to 0.5, in small swarms we find values between 0.5 and 1, being close to 1 for the smallest swarms (about 10 individuals).

This does not mean that mosquitoes in small swarms explore larger regions; rather, it indicates that swarms themselves occupy a narrower area. As a result, localization cannot be properly detected. When we consider both small and large swarms together, see Fig.S1f,g, we find that both ratios reach the plateau of 0.5 when the number of individuals is approximately equal to 100.

**Speed localization.** In the main text we showed that fluctuations of individual mean speeds cannot be explained by statistical fluctuations alone. This was done by comparing the standard error of the speed (over all mosquitoes at all instants in times), $\Sigma$, with the standard deviation of individual mean speeds, $\sigma_S$. In small swarms, as in large ones, we find that $\sigma_S$ is greater than $\Sigma$. But, the separation between these two quantities is less pronounced that in the case of large swarms, see Fig.S1c,h. Two main factors contribute: first in small groups, we expect a higher standard error, due to the definition of $\Sigma$, see Methods. Second, in small swarms we find higher overall variability. As a consequence, working only on small swarms, we would not be able to clearly discriminate between statistical fluctuations and localization.

**Spatial correlation function of the speed fluctuations.** This is by far the most relevant difference between small and large swarms. In small swarms, the connected correlation function of the speed does not present a characteristic trend with the distance. Depending on the swarm, the correlation can be positive for short distances and then it decreases until it crosses the $x$-axis, as for the light green plot of Fig.S1d; it can also be positive for very short distances before abruptly becoming negative, or even be negative from the outset, as in the dark green plot. This behavior is not simply due to the spatial size of the swarm, as we show in Fig.S1d, where the three correlation functions shown are all from swarms of the same size, $L = 0.8$m. In contrast, in large swarms, we observe a typical trend, see the purple plots in Fig.S1i: regardless the size of the swarm, the correlation starts positive, and then it smoothly decreases to cross the $x$-axis.

The linear dependence between the correlation length, $\xi$, and the size of the system, $L$, which indicates the scale–free nature of the correlation, is not evident in small swarms, see Fig.S1e. The Pearson's coefficient drops from 0.85 in large swarms to 0.61 in small swarms (with a p–value smaller than $10^{-6}$ in large swarm and equal to $10^{-6}$ in small ones), suggesting that there is not strong linear dependence between the two quantities. When we consider the full dataset (large and small swarms combined) on the same plot, Fig.S1k, the linear trend re-emerges, with Pearson's coefficient returning to $\rho = 0.82$ (only 0.03 lower than that obtained in the main text for the large-swarm dataset alone) indicating that small swarms contribute little to the linear dependence. Moreover, in 17 out of 41 small swarms (in the 41% of the swarms), the correlation length $\xi$ is equal to zero, meaning that even for very short distances (for $r$ between 0 and the nearest-neighbor distance) the correlation is negative; this is a phenomenon that we never find in large swarms.

This analysis confirms that using data on small swarms is not suitable for the study of collective behavior. With a dataset of only small swarms, we would neither be able to observe spatial and speed localization, nor scale–free correlation. Furthermore, small swarms do not provide useful additional information to large swarms for this kind of behavioral analysis. This does not mean that small swarms cannot provide useful information at a general level, for instance from a mechanosensory point of view, but only that they are not suitable to study interaction rules generating collective behavior.

#### 2 Spatial correlation function of the velocity fluctuations

Following previous studies on insect swarms [3, 4], we computed the connected form of the spatial correlation function of the velocity vectors.

To this aim, at each instant in time, we computed the average velocity of the swarm, $\vec{V}(t)$, as:

$$\vec{V}(t) = \frac{1}{N}\sum_{i=1}^{N} \vec{v}_i(t) \tag{1}$$

where $N$ is the number of mosquitoes and $\vec{v}_i(t)$ the velocity vector of mosquito $i$ at time $t$.

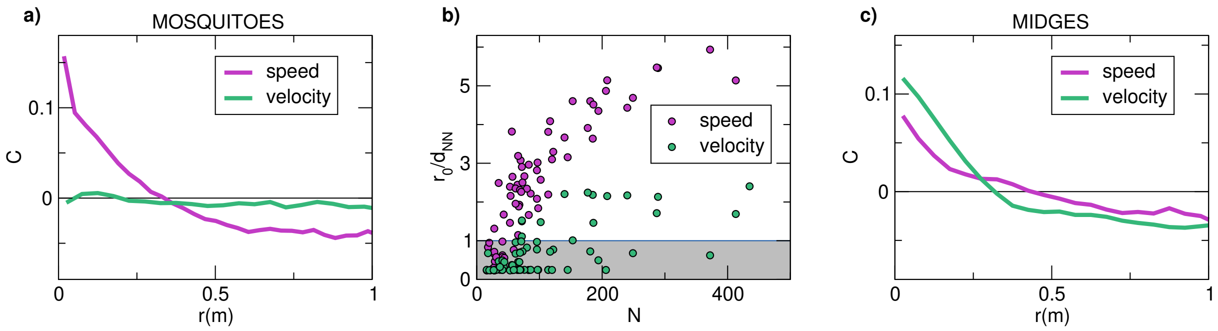


**Fig. S2 Spatial correlation function of the velocity fluctuations. a.** Spatial correlation function of the speed and of the velocity for the swarm 20200928_$ACQ3$. In purple the correlation function of the speed, already presented in the main text. In green the correlation function of the velocity. Unlike other systems, our swarms shows no significant correlation of the velocity. The function does not have a particular trend for small $r$, while it is almost equal to zero at all distances. **b.** The correlation range measured with respect to the typical distance to the nearest neighbor, $d_{NN}$, as a function of the number of mosquitoes in the swarm, $N$. The purple circles refer to the correlation range of the speed, showing that this distance increases with the size of the swarm, reaching a plateau at around $6d_{NN}$. For most of the swarm, instead, the correlation range of the velocity is smaller than $d_{NN}$ (light gray region), confirming that the correlation is not significant. **c.** Spatial correlation function of the speed (purple) and of the velocity (green) for a swarm of midges (*Diptera Chironomidae*). The correlation function of the speed and that of the velocity show a similar trend, with the velocity correlation being dominant. Data for midges correlation belong to CoBBS (Collective Behavior in Biological System) group.

We used $\vec{V}(t)$ to compute individual velocity fluctuations, $\delta\vec{v}_i(t)$, for each mosquito $i$ at any given time $t$:

$$\delta\vec{v}_i(t) = \vec{v}_i(t) - \vec{V}(t) \tag{2}$$

With the same approach we used for the correlation of the speed (see Methods from the main text), we computed the connected correlation function of the velocity at time $t$:

$$\hat{C}(r,t) = \frac{\sum_{i,j} \delta\vec{v}_i(t) \cdot \delta\vec{v}_j(t)\delta(r_{ij}(t), r)}{\sum_{i,j} \delta(r_{ij}(t), r)} \tag{3}$$

where $r_{ij}$ is the distance between mosquito $i$ and mosquito $j$, the $\cdot$ indicates the scalar product, and $\delta(r_{ij}(t), r)$ denotes the Kronecker delta, which is equal to 1 when $r_{ij}(t) = 1$ and 0 otherwise.

Finally, we normalized for $\hat{C}(0,t)$ and averaged over time. So that we define the correlation function as:

$$C(r) = \frac{1}{T}\sum_{t=0}^{T} \frac{\hat{C}(r,t)}{C(0,t)} \tag{4}$$

where $T$ is the duration of the acquisition.

In Fig.S2a, we show the comparison between the spatial correlation of the speed and of the velocity. Unlike the correlation of the speed, the correlation of the velocity displays very low values at all the distances. It does not have the typical trend of the speed correlation. Its correlation range, $r_0$, computed as the first distance $r$ such that the correlation function crosses the 0, is small in all the 71 swarms of the dataset. The ratio between the correlation range and the distance from the nearest neighbor, $r_0/d_{NN}$, is in most of the cases smaller than 1 (see Fig.S2b), indicating that the correlation is not significant. As a comparison, we show in Fig.S2c the two correlation functions for a swarm of midges, where in both cases we find a non–trivial trend.

### 3 Lack of spatial trend in speed.

We investigate potential trends in mosquito speed fluctuations as a function of their distance from the center of the swarm, $d_{3D}$, and their position in three dimensional space, namely the three spatial coordinates $x$, $y$, and $z$. The joint probability distribution of speed fluctuations with respect to the positional quantities $d_{3D}$, $x$, $y$ and $z$, show that speed fluctuations are mostly centered around 0m/s regardless of mosquito position within the swarm, see Fig.S3. Superimposed scatter plots of speed fluctuations as a function of position variables further show that there are no specific

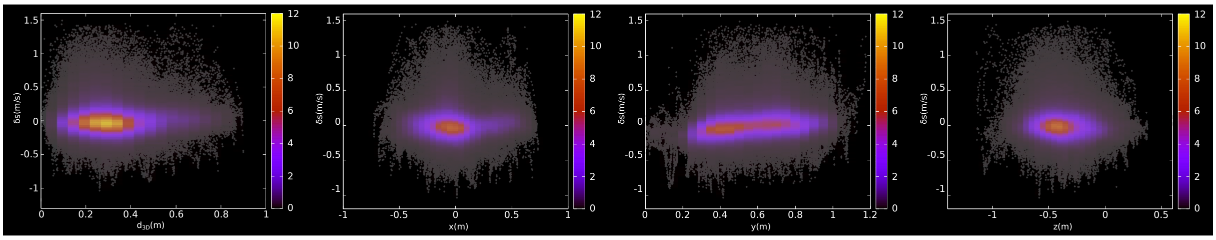

**Fig. S3 Lack of spatial trend of the speed.** The joint probability distribution of speed fluctuations with respect to the distance from the center of the swarm $d_{3D}$ (panel **a**), and the three spatial coordinates $x$ (panel **b**), $y$ (panel **c**) and $z$ (panel **d**), show that speed fluctuations are mostly centered around 0m/s regardless of mosquito position within the swarm. Superimposed scatter plots of speed fluctuations (highlighted in gray) as a function of position variables further show that there are no specific regions where mosquitoes tend to fly faster or slower than the group average, with large fluctuations uniformly spread throughout the swarm volume.

regions where mosquitoes tend to fly faster or slower than the group average, with large fluctuations uniformly spread throughout the swarm volume.

| DATE | ACQ | N | $R(m)$ | $H(m)$ | $r(m)$ | $h(m)$ | $\sigma_S(m/s)$ | $\Sigma(m/s)$ | $r_0(m)$ | $\xi(m)$ | $L(m)$ |
|---|---|---|---|---|---|---|---|---|---|---|---|
| 20200923 | 3 | 113 | 0.21 | 0.13 | 0.12 | 0.05 | 0.059 | 0.015 | 0.27 | 0.09 | 0.98 |
| | 4 | 95 | 0.20 | 0.11 | 0.12 | 0.05 | 0.056 | 0.016 | 0.20 | 0.06 | 0.95 |
| 20200928 | 1 | 183 | 0.24 | 0.13 | 0.14 | 0.05 | 0.089 | 0.016 | 0.27 | 0.10 | 1.52 |
| | 2 | 209 | 0.23 | 0.15 | 0.13 | 0.05 | 0.067 | 0.014 | 0.33 | 0.11 | 1.41 |
| | 3 | 206 | 0.24 | 0.16 | 0.13 | 0.05 | 0.063 | 0.012 | 0.34 | 0.11 | 1.29 |
| 20210413 | 4 | 96 | 0.20 | 0.12 | 0.12 | 0.05 | 0.130 | 0.020 | 0.17 | 0.06 | 1.32 |
| 20210525 | 4 | 102 | 0.19 | 0.11 | 0.13 | 0.06 | 0.082 | 0.021 | 0.20 | 0.07 | 1.06 |
| | 5 | 175 | 0.21 | 0.13 | 0.13 | 0.06 | 0.072 | 0.015 | 0.28 | 0.09 | 1.17 |
| | 6 | 139 | 0.25 | 0.15 | 0.13 | 0.05 | 0.073 | 0.016 | 0.30 | 0.11 | 1.24 |
| 20241209 | 2 | 82 | 0.20 | 0.10 | 0.14 | 0.05 | 0.072 | 0.019 | 0.14 | 0.04 | 1.07 |
| | 4 | 118 | 0.34 | 0.12 | 0.20 | 0.07 | 0.086 | 0.015 | 0.31 | 0.11 | 1.64 |
| | 6 | 146 | 0.39 | 0.13 | 0.21 | 0.07 | 0.069 | 0.013 | 0.31 | 0.11 | 1.72 |
| | 10 | 122 | 0.39 | 0.12 | 0.15 | 0.06 | 0.069 | 0.015 | 0.33 | 0.10 | 1.51 |
| | 11 | 181 | 0.42 | 0.15 | 0.23 | 0.08 | 0.081 | 0.013 | 0.49 | 0.16 | 2.00 |
| | 12 | 153 | 0.41 | 0.11 | 0.16 | 0.06 | 0.065 | 0.013 | 0.44 | 0.12 | 1.67 |
| | 14 | 114 | 0.40 | 0.13 | 0.16 | 0.06 | 0.059 | 0.014 | 0.23 | 0.08 | 1.57 |
| | 15 | 116 | 0.39 | 0.14 | 0.17 | 0.07 | 0.065 | 0.014 | 0.43 | 0.15 | 1.60 |
| 20241216 | 3 | 434 | 0.51 | 0.19 | 0.17 | 0.07 | 0.083 | 0.009 | 0.57 | 0.17 | 2.16 |
| | 4 | 288 | 0.45 | 0.19 | 0.17 | 0.07 | 0.082 | 0.011 | 0.51 | 0.16 | 2.02 |
| | 5 | 249 | 0.43 | 0.18 | 0.18 | 0.07 | 0.084 | 0.011 | 0.46 | 0.14 | 1.94 |
| | 6 | 187 | 0.39 | 0.19 | 0.17 | 0.07 | 0.079 | 0.012 | 0.45 | 0.16 | 1.86 |
| | 7 | 410 | 0.44 | 0.16 | 0.22 | 0.08 | 0.096 | 0.011 | 0.45 | 0.13 | 2.24 |
| | 8 | 289 | 0.45 | 0.16 | 0.18 | 0.08 | 0.085 | 0.011 | 0.52 | 0.16 | 2.02 |
| | 9 | 193 | 0.43 | 0.16 | 0.19 | 0.08 | 0.078 | 0.011 | 0.46 | 0.16 | 1.95 |
| | 10 | 373 | 0.42 | 0.14 | 0.18 | 0.07 | 0.083 | 0.008 | 0.50 | 0.17 | 2.10 |
| | 11 | 241 | 0.40 | 0.13 | 0.20 | 0.08 | 0.091 | 0.013 | 0.39 | 0.12 | 1.98 |
| 20241217 | 3 | 98 | 0.28 | 0.13 | 0.17 | 0.06 | 0.074 | 0.018 | 0.17 | 0.08 | 1.31 |
| | 5 | 97 | 0.28 | 0.14 | 0.17 | 0.06 | 0.079 | 0.018 | 0.27 | 0.10 | 1.32 |
| | 6 | 83 | 0.24 | 0.12 | 0.16 | 0.06 | 0.063 | 0.016 | 0.27 | 0.09 | 1.13 |
| | 8 | 85 | 0.26 | 0.12 | 0.18 | 0.06 | 0.108 | 0.020 | 0.22 | 0.07 | 1.30 |

**Table 1 Summary of the data.** DATE and ACQ identify the date of the data collection and the number of the acquisition. On the same day we often perform several acquisitions. Data belong to two different experimental campaigns. In $2020-2021$ data are collected at 170fps and each acquisition last $\sim$ 15.9s, in 2024 at 80fps and last $\sim$ 33s. Time duration is limited by hardware constraints. $N$ is the number of mosquitoes in the swarm. $R$(m) is the radius of the swarm on the horizontal plane, computed as the mean value of the group spatial distribution. $H$(m) is the elongation of the swarm, i.e. its vertical extent computed as the mean value of the spatial distribution on the $y$-axis. $r$(m) is the typical radius of individual trajectories on the horizontal plane, computed averaging, over all mosquitoes, the mean values of individual spatial distributions. $h$(m) is the typical elongation of individual trajectories on the horizontal plane, computed averaging, over all mosquitoes, the mean values of individual spatial distributions. $\sigma_S$(m/s) standard deviation of the mean individual speed. $\Sigma$(m/s) is the standard error of the distribution of the fluctuations of the speed. $r_0$(m) is the correlation range. $\xi$(m) is the correlation length. $L(m)$ is the size of the swarm.

#### References

[1] Anderson, P.W.: More is different. Science **177**(4047), 393–396 (1972)

[2] Cavagna, A., Giardina, I., Gucciardino, M.A., Iacomelli, G., Lombardi, M., Melillo, S., Monacchia, G., Parisi, L., Peirce, M.J., Spaccapelo, R.: Characterization of lab-based swarms of anopheles gambiae mosquitoes using 3d-video tracking. Scientific Reports **13** (2023)

[3] Attanasi, A., Cavagna, A., Del Castello, L., Giardina, I., Melillo, S., Parisi, L., Pohl, O., Rossaro, B., Shen, E., Silvestri, E., Viale, M.: Collective behaviour without collective order in wild swarms of midges. PLoS Comput Biol **10(7)** (2014)

[4] Kelley, D., Ouellette, N.: Emergent dynamics of laboratory insect swarms. Scientific reports **3**, 1073 (2013)
